# Orb-dependent polyadenylation contributes to PLP expression and centrosome scaffold assembly

**DOI:** 10.1101/2021.10.29.466388

**Authors:** Junnan Fang, Dorothy A. Lerit

**Author notes:** Correspondence: Dorothy A. Lerit, Ph.D.

## Abstract

As the microtubule-organizing centers (MTOCs) of most cells, centrosomes engineer the bipolar mitotic spindle required for error-free mitosis. *Drosophila* Pericentrin (PCNT)-like protein (PLP) is a key centrosome component that directs formation of a pericentriolar material (PCM) scaffold required for PCM organization and MTOC function. Here, we investigate the post-transcriptional regulation of *plp* mRNA. We identify conserved binding sites for cytoplasmic polyadenylation element binding (CPEB) proteins within the *plp* 3’-untranslated region and examine the role of the CPEB ortholog, oo18 RNA-binding protein (Orb), in *plp* mRNA regulation. Our data show Orb biochemically interacts with *plp* mRNA and promotes PLP protein expression. Loss of *orb*, but not *orb2*, diminishes PLP levels in embryonic extracts. Consequently, PLP localization to centrosomes and function in PCM scaffolding is compromised in *orb* mutant embryos, resulting in genome instability and embryonic lethality. Moreover, we find PLP over-expression can restore centrosome scaffolding and rescue the cell division defects caused by *orb* depletion. Our data suggest Orb modulates PLP expression at the level of *plp* mRNA polyadenylation and showcases the post-transcriptional regulation of core, conserved centrosomal mRNAs as critical for centrosome function.

## Introduction

Centrosomes are microtubule-organizing centers that function in spindle assembly during cell division, cilia and flagella formation, and intracellular trafficking (Vertii et al., 2016). The centrosome comprises a pair of centrioles embedded in pericentriolar material (PCM), a matrix of proteins that directs microtubule nucleation and organization (Palazzo et al., 1999). Centrosome function relies on cell cycle-dependent oscillations in PCM recruitment before mitotic onset, followed by PCM shedding at mitotic exit (Gould and Borisy, 1977; Khodjakov and Rieder, 1999). The mechanisms that regulate oscillations in centrosome activity remain incompletely understood.

In humans, PCM recruitment and microtubule organization is supported by Pericentrin (PCNT) (Dictenberg et al., 1998; Zimmerman et al., 2004; Haren et al., 2009; Lee and Rhee, 2011; Chen et al., 2014; Kim et al., 2015). Consequently, deregulation of *PCNT* results in severe human genetic disorders, such as Trisomy 21/Down syndrome and microcephalic osteodysplastic primordial dwarfism type II (MOPD II) (Jurczyk et al., 2004; Rauch et al., 2008; Delaval and Doxsey, 2010; Galati et al., 2018). Understanding how *PCNT* expression is regulated in healthy tissues and deregulated in developmental disorders remains a critical challenge.

In *Drosophila*, PCNT-like protein (PLP) is the ortholog of *PCNT* (Martinez-Campos et al., 2004) with conserved roles in PCM recruitment and scaffolding and microtubule organization (Dobbelaere et al., 2008; Lerit and Rusan, 2013; Galletta et al., 2014; Lerit et al., 2015; Richens et al., 2015; Roque et al., 2018; Galletta et al., 2020). Additionally, the spatial configuration of PCNT and PLP at centrosomes is identical, with the C-termini proximal to centrioles and the N-termini extended out into the PCM (Fu and Glover David; Lawo et al., 2012; Mennella et al., 2012; Sonnen et al., 2012). Intriguingly, aspects of *PCNT* and *plp* mRNA post-transcriptional regulation are also conserved, as both *PCNT* and *plp* mRNAs localize to centrosomes (Lécuyer et al., 2007; Sepulveda et al., 2018; Chouaib et al., 2020; Ryder et al., 2020; Safieddine et al., 2021). Therefore, *Drosophila* PLP is a valuable model to study centrosome regulation likely to inform mechanisms of human disease linked to *PCNT* dysfunction.

How and why *PCNT* or *plp* mRNAs localize to the centrosome are largely unknown, although recent work implicates a co-translational transport mechanism (Sepulveda et al., 2018; Chouaib et al., 2020; Safieddine et al., 2021). RNA localization coupled with translational control is a conserved regulatory paradigm used to generate spatial enrichments in gene activity and critical for diverse cellular processes, reviewed in (Cody et al., 2013; Buxbaum et al., 2014; Ryder and Lerit, 2018). RNA localization and translational control are often regulated by RNA-binding proteins (RBPs) recognizing cis-elements within the 3’-untranslated regions (UTRs) of target RNAs (Kislauskis et al., 1994).

One conserved family of RBPs implicated in RNA localization and translational control is the cytoplasmic polyadenylation element binding (CPEB) proteins (Mendez and Richter, 2001). CPEB binds to target mRNAs through recognition of cytoplasmic polyadenylation element (CPE) motifs and promotes mRNA polyadenylation following phosphorylation by Aurora A kinase (Mendez et al., 2000a; Mendez et al., 2000b; Hodgman et al., 2001). While CPEB promotes the translation of some mRNA targets, such as *c-mos*, *p53*, *Cyclin B1*, and *casein* mRNAs (Stebbins-Boaz et al., 1996; Groisman et al., 2000; Mendez et al., 2000b; Cao and Richter, 2002; Choi et al., 2004; Burns and Richter, 2008; Burns et al., 2011), it represses the translation of others, including *myc* mRNA (Groisman et al., 2006). Further, CPEB can mediate localization of its target mRNAs independent of its polyadenylation or translation activities, as shown with *zonal occludens-1* mRNA (Nagaoka et al., 2012).

*Drosophila* encodes two CPEB proteins, oo18 RNA-binding protein (Orb) and Orb2 (Lantz et al., 1992; Hafer et al., 2011). While Orb2 functions are best defined in the testis and central nervous system (Keleman et al., 2007; Mastushita-Sakai et al., 2010; Hafer et al., 2011; Xu et al., 2012), Orb regulates RNA localization and translation during oogenesis and early embryogenesis (Lantz et al., 1992; Christerson and McKearin, 1994; Chang et al., 2001; Castagnetti and Ephrussi, 2003; Rojas-Ríos et al., 2015). Orb is orthologous to mammalian CPEB1 and similarly promotes polyadenylation to regulate the stability and translation of its target mRNAs, including *oskar* (*osk*), *gurken* (*grk*), and *Autophagy-related 12* (*Atg12*) mRNAs (Christerson and McKearin, 1994; Chang et al., 1999; Chang et al., 2001; Castagnetti and Ephrussi, 2003; Norvell et al., 2015; Rojas-Ríos et al., 2015; Davidson et al., 2016).

Recent transcriptomics identified numerous mRNA targets of Orb from *Drosophila* S2 cells and CPEB1 in cultured mammalian cells (Stepien et al., 2016; Pascual et al., 2021). Among the Orb/CPEB1 mRNA targets are several mRNAs encoding centrosome proteins. In *Xenopus* and cultured mammalian cells, CPEB proteins also localize to centrosomes, and mRNAs localizing to spindle poles show enrichment of CPE motifs in their 3’-UTRs (Groisman et al., 2000; Blower et al., 2007; Eliscovich et al., 2008; Sharp et al., 2011; Pascual et al., 2021). These data are consistent with a model whereby CPEB proteins regulate components of the mitotic machinery. However, whether Orb contributes to the post-transcriptional regulation of centrosomal mRNAs remains to be investigated.

In this study, we analyzed the common mRNA targets of human CPEB1 and *Drosophila* Orb and identified *PCNT*/*plp* mRNA. To test a role for Orb in regulating *plp* mRNA, we identified a specific biochemical association between Orb and *plp* mRNA and examined the role of Orb in *plp* mRNA localization to centrosomes and translational control. Although dispensable for *plp* mRNA localization, Orb specifically regulates PLP protein expression by promoting polyadenylation of the short *plp* 3’-UTR. We demonstrate Orb contributes to PLP protein localization and PCM organization at centrosomes, required for genome stability. Our data suggest *plp* mRNA is a critical target of Orb required for embryonic viability and highlight the translational regulation of centrosomal mRNAs as important for centrosome activity and function.

## Results

### *plp* mRNA localizes at centrosomes during early embryogenesis

During the first two hours of development, the *Drosophila* embryo develops as a syncytium whereby the somatic nuclei rapidly divide through abridged (S-M) mitotic cycles and migrate to the embryonic cortex prior to cellularization during nuclear cycle (NC) 14 (Rothwell and Sullivan, 1999). We recently reported that multiple mRNAs encoding proteins required for centrosome function, including *plp* mRNA, localize to embryonic centrosomes in a cell cycle-dependent manner (Ryder et al., 2020). To further analyze how *plp* mRNA localization to the centrosome is regulated by cell cycle progression, we performed single molecule fluorescence in situ hybridization (smFISH) for *plp* mRNA and a control transcript, *gapdh* mRNA, throughout syncytial development and quantified mRNA distributions relative to centrosomes labeled with GFP-γ-tubulin (GFP-γTub). Despite relatively fewer smFISH signals, consistent with lower levels of *plp* mRNA expression relative to *gapdh* (Graveley et al., 2011), some molecules of *plp* mRNA overlap with the centrosomes. In contrast, *gapdh* mRNA disperses throughout the cytoplasm (**Figure 1A, B**).

**Figure 1.**
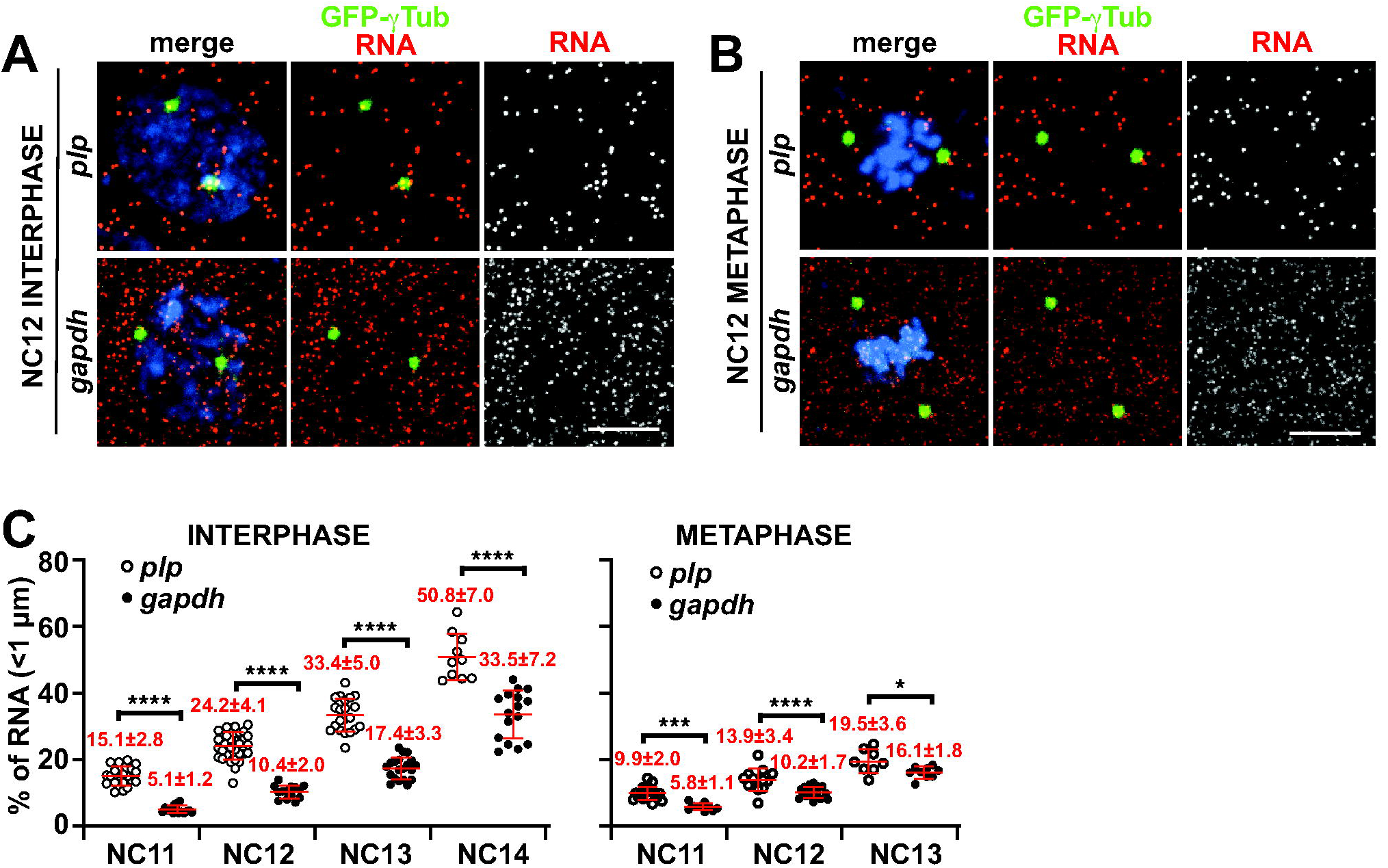
*plp* mRNA localizes to centrosomes. Maximum intensity projections of NC 12 control embryos expressing GFP-γTub (green) labeled with *plp* or *gapdh* smFISH probes (red) in (A) interphase and (B) metaphase. (C) Scatter dot plots show the quantification of *plp* vs. *gapdh* mRNA enrichment within 1 μm from the centrosome surface at different NCs. Supplemental Table 1 lists the number of embryos, centrosomes, and RNA objects quantified per condition. Mean ± S.D. are displayed. n.s, not significant; \**p*<0.05; \*\*\**p*<0.001; \*\*\*\**p*<0.0001 by one-way ANOVA followed by Dunnett’s T3 multiple comparisons test. Bars: 5 μm.

To quantitatively compare the distributions of *plp* versus *gapdh* mRNAs, we measured the percent of mRNA residing within 1 μm from the centrosome surface using a custom Python package (Ryder et al., 2020; Ryder and Lerit, 2020). From this analysis, we observed significantly more *plp* mRNA localized to centrosomes relative to *gapdh* mRNA (2∼3-fold more *plp* mRNA than *gapdh* mRNA at interphase versus 1.5∼2-fold more at metaphase centrosomes; **Figure 1C, Table S1**). As the syncytial embryo undergoes successive nuclear divisions, the size of each pseudo-cell becomes progressively smaller (Foe and Alberts, 1983). Consequently, the percentage of mRNA residing within 1 μm from the centrosome surface increases with each NC stage. Nonetheless, *plp* mRNA remains significantly enriched at centrosomes relative to *gapdh* throughout syncytial development, particularly during interphase. Furthermore, imaging of *plp* mRNA together with a *GFP-PLP* transgene reveals *plp* mRNA and protein partially overlap at centrosomes, as recently detected in cultured mammalian cells (Sepulveda et al., 2018; Safieddine et al., 2021) (**Figure S1A, B**). Taken together, we conclude that *plp* mRNA localizes to centrosomes, likely via a dynamic process that may involve co-translational transport.

### *plp* mRNA associates with Orb protein

RNA localization coupled with translational control is a conserved mechanism that functions in various cellular contexts and is commonly regulated by RBPs, reviewed in (Buxbaum et al., 2014; Das et al., 2021). Transcriptomics uncovered thousands of direct mRNA targets of Orb and Orb2 in *Drosophila* S2 cells (Stepien et al., 2016). A similar analysis in HeLa cells also uncovered thousands of mRNA targets of orthologous CPEB1 and CPEB4 (Pascual et al., 2021). We compared the RNA substrates of homologous Orb and CPEB1 proteins identified in the Stepien and Pascual datasets and found 195 common genes (Hake and Richter, 1994)(**Figure 2A; Table S2**). Analysis of the gene ontologies (GO) within the common genes using the cellular component function in PANTHER uncovered significantly enriched ontologies related to the centrosome, as represented by *PCNT/plp, CEP192/Spd-2,* and *PLK1/Polo* mRNAs (**Figure S2A, Table S2**) (Mi et al., 2021). In accordance, recent work shows CPEB1 is required for the translation and recruitment of PLK1 to centrosomes (Pascual et al., 2021). Here, we focused our subsequent analysis on PLP.

**Figure 2.**
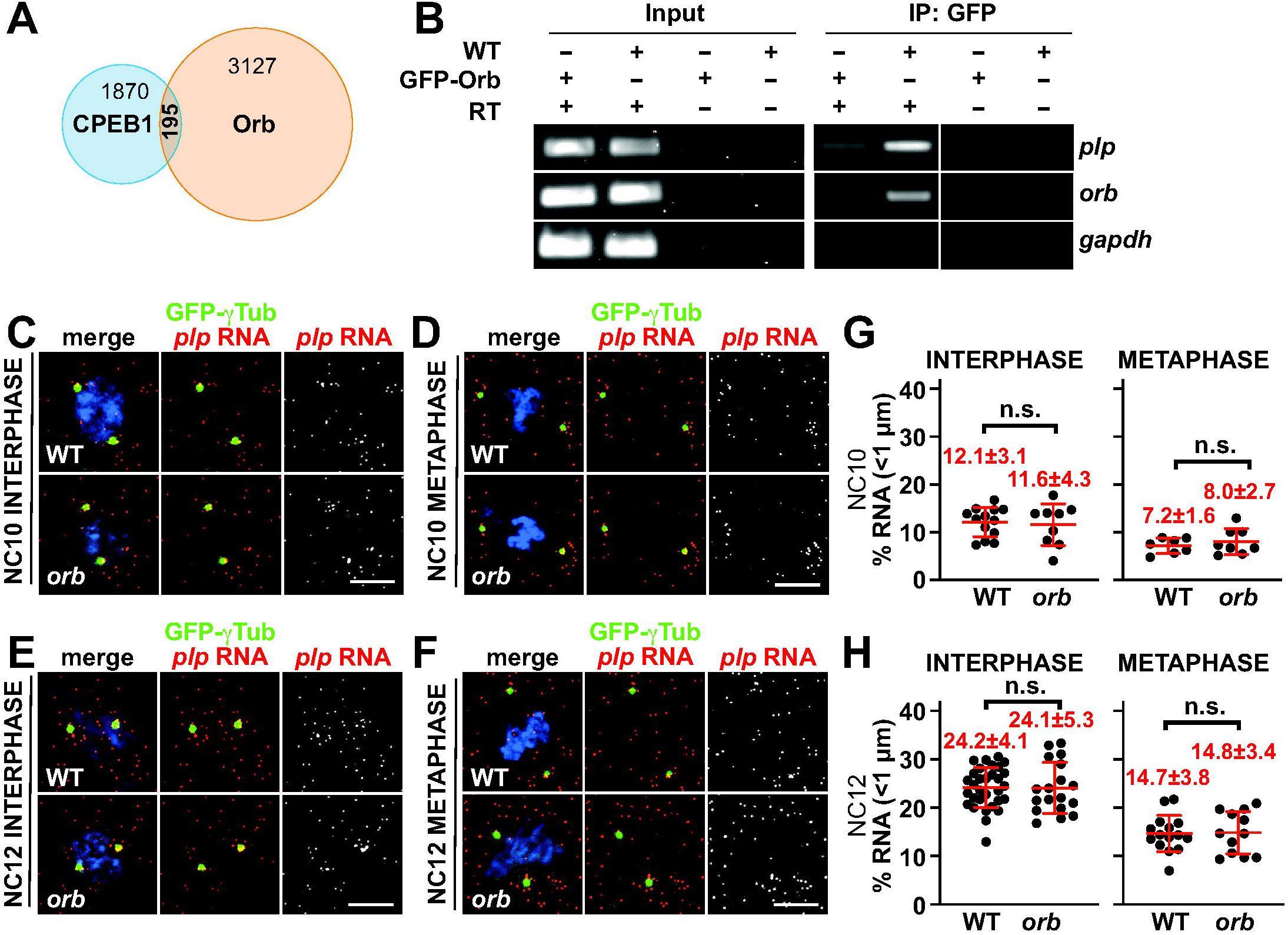
Orb interacts with *plp* mRNA, but is dispensable for *plp* mRNA localization. (A) Venn diagram shows the common mRNA targets of CPEB1 and Orb (Stepien et al., 2016; Pascual et al., 2021). Supplemental Table 2 lists overlapping mRNA targets. (B) Gels show RNA-immunoprecipitation results from WT vs. *GFP-Orb* ovarian extracts, as performed with anti-GFP beads. Input shows 10% of the total RNA. cDNAs were amplified with *plp, orb*, or *gapdh* primers. *orb* is a positive control (Tan et al., 2001), while *gapdh* serves as a negative control. Images show maximum intensity projections of control and *orb^F343^*/*orb^mel^* mutant (C-D) NC 10 and (E-F) NC 12 embryos expressing GFP-γTub (green) and labeled with *plp* smFISH probes (red). (G-H) Scatter plots show the quantification of *plp* mRNA localization within 1 μm from the centrosome surface in NC 10 or NC 12 embryos of the indicated genotypes. Supplemental Table 1 lists the number of embryos, centrosomes, and RNA objects quantified per condition. Mean ± S.D. are displayed. n.s, not significant by unpaired t-test. Bars: 5 μm. Uncropped gels are available to view on Figshare: https://doi.org/10.6084/m9.figshare.16900417.v1

Most Orb targets contain a CPE consensus motif (e.g., UUUUAU or UUUUAAU) in their 3’-UTR (Fox et al., 1989; McGrew and Richter, 1990), although CPEB proteins can also recognize non-canonical CPE motifs (e.g., UUUUACU or UUUUAAGU (Piqué et al., 2008) or UUUUAA (Barkoff et al., 2000) or UUUUGU (Stepien et al., 2016)). As a first step of validating *plp* mRNA as an Orb target, we identified two consensus CPE motifs within the *plp* 3’UTR (Figure S2B, red boxes). Next, we aligned the *plp* 3’-UTR across multiple *Drosophila* species using the conservation insect track on the UCSC genome browser and found these CPE motifs were conserved across millions of years of evolutionary distance (data not shown)(Kent et al., 2002). Finally, we aligned *Drosophila melanogaster plp* and human *PCNT* 3’UTRs using Clustal Omega (Madeira et al., 2019). *plp* transcripts utilize one of two different 3’-UTRs varying in length: a short 3’-UTR (431 nt) or a long 3’-UTR (591 nt), as shown in **Figure S2B** (Graveley et al., 2011). The short *plp* 3’-UTR has 100% identity to the long *plp* 3’-UTR and 44.6% identity to the *PCNT* 3’-UTR, while the long *plp* 3’-UTR has 46.9% identity to the *PCNT* 3’-UTR. All three 3’-UTRs contain 2 consensus CPE motifs (**Figure S2B,** red boxes). Conservation of CPE sites within the *plp* 3’UTR argues they are likely to be biologically relevant.

To investigate whether Orb associates with *plp* mRNA during early development, we used a GFP-Orb gene-trap, which inserts *gfp* coding sequences at the endogenous *orb* locus (Nagarkar-Jaiswal et al., 2015), to affinity purify interacting mRNAs by RT-PCR. Similar to endogenous *orb* expression, we detected GFP-Orb more readily in ovaries as compared to embryos (Lantz et al., 1992; Lantz et al., 1994; Rangan et al., 2009; Hafer et al., 2011) (**Figure S3A,B**). Using *GFP-Orb* ovarian extracts, we confirmed Orb associates with *orb* mRNA, consistent with prior work (Tan et al., 2001). We also detected a specific interaction with *plp* mRNA, but not the negative control, *gapdh* mRNA (**Figure 2B**). We conclude *plp* mRNA interacts with Orb during early development, suggesting Orb may contribute to the post-transcriptional regulation of *plp* mRNA.

### Orb is dispensable for *plp* mRNA localization

To examine if Orb contributes to *plp* mRNA localization to centrosomes, we compared *plp* mRNA distributions using smFISH in control versus *orb* mutant embryos. To examine maternal effects, we harvested embryos from transheterozygous *orb^F343^*/*orb^mel^* (hereafter, *orb*) mutant mothers. While *orb^F343^* is a protein null, *orb^mel^* is a weak hypomorph (Christerson and McKearin, 1994; Lantz et al., 1994). By western blot, we confirmed Orb protein expression is reduced in *orb* mutant ovarian extracts, consistent with prior work (**Figure S3C**) (Christerson and McKearin, 1994; Lantz et al., 1994). At all stages examined, we observed no difference in *plp* mRNA localization in controls versus *orb* mutant embryos (**Figure 2C–H**). *gapdh* mRNA was also unaffected (**Figure S4**). These results indicate that Orb is not required for *plp* mRNA localization to centrosomes.

### Orb promotes PLP translation

Next, to investigate the contribution of Orb in regulating PLP protein expression, we performed semi-quantitative western blotting to examine PLP levels in 0-2 hr wild-type (WT) and *orb* mutant embryos. The *plp* gene encodes 12 different protein isoforms (Graveley et al., 2011). Consistently, we observe multiple PLP isoforms within embryonic extracts (**Figure 3A-D**), similar to prior observations using larval brain extracts (Galletta et al., 2014).The longest PLP isoforms are predicted to migrate ∼320 KDa; therefore, we quantified these bands to analyze PLP levels in various genetic backgrounds (**Figure 3A-D**). In *orb* mutant embryos, we observed PLP protein is ∼50% downregulated (**Figure 3A, E**). We recently showed *plp* hemizygote embryos (*plp^2172^*/+) also result in an average ∼50% downregulation of PLP protein (Fang and Lerit, 2020). Therefore, we generated *plp^2172^*, *orb^F343^* recombinant chromosomes and crossed these animals to *orb^mel^* mutants to harvest embryos from *plp* hemizygous, *orb* mutant mothers (genotype: *plp^2172^*/+, *orb^F343^*/*orb^mel^*; hereafter, *plp*/+, *orb*). Quantification shows simultaneous depletion of *plp* and *orb* within *plp*/+, *orb* mutant embryos results in ∼75% reduction of PLP protein, further demonstrating a requirement of *orb* for PLP translation (**Figure 3B, E**).

**Figure 3.**
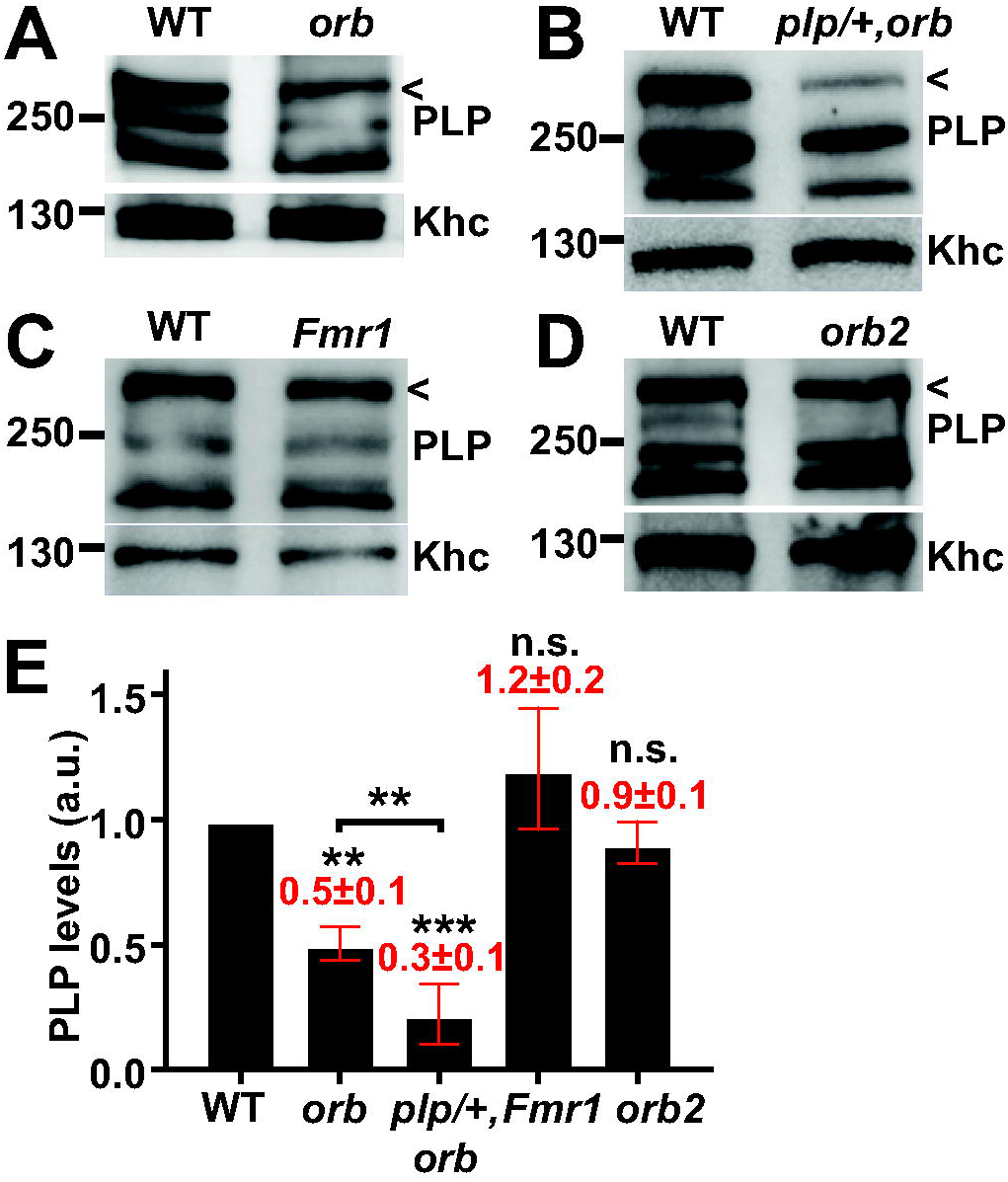
Orb specifically promotes PLP protein expression. (A-D) Immunoblots show PLP levels relative to the Khc (anti-SUK4) load control in 0-2 h *Drosophila* embryo extracts from the indicated genotypes, *orb* mutant: *orb^F343^*/*orb^mel^*; *plp*/+,*orb* mutant: *plp^2172^*/+, *orb^F343^*/*orb^mel^*. The upper PLP band (carrot) was used for quantification. (E) Quantification of relative PLP levels normalized to WT. Significance was determined by one-way ANOVA followed by Dunnett’s T3 multiple comparisons test relative to WT; bracket marks unpaired t-test. Mean ± S.D. are displayed. n.s, not significant; \*\**p*<0.01; \*\*\**p*<0.001. Uncropped blots are available to view on Figshare: https://doi.org/10.6084/m9.figshare.16900423.v1

To investigate the specificity of the *orb*-dependent response on PLP expression, we also examined PLP levels after depleting the other *Drosophila* CPEB family member, Orb2, and Fragile X Mental Retardation Protein (FMRP), an RNA-binding protein known to share several common mRNA targets with *orb* (Costa et al., 2005). We note no significant difference in PLP levels in embryos derived from *orb2* or *Fmr1* null mutant mothers (**Figure 3C–E**). These results highlight a specific requirement of Orb to support normal levels of PLP protein, likely at the level of translational activation.

### Orb regulates PLP-Cnn scaffold organization

In interphase embryos, PLP normally localizes to the centriole and at the tips of extended PCM flares (Lerit et al., 2015; Richens et al., 2015). To investigate whether the depletion of PLP protein observed in *orb* mutants alters PLP localization to the centrosome, we quantified the intensity of endogenous PLP signals at control versus *orb* mutant centrosomes in NC 13 embryos (**Figure 4**). As expected, WT embryos had PLP signals at the centriole and flare zones (**Figure 4A,** WT). We generated *plp* null mutant germline clone embryos (*plp* mutants) using the FLP/*ovo*^D^ method (Chou and Perrimon, 1996) and confirmed PLP is undetectable, validating the specificity of our antibody (**Figure 4A,** *plp*). In contrast, *orb* and recombinant *plp*/+, *orb* mutant embryos show reduced PLP localization to centrosomes compared to WT centrosomes (**Figure 4A;** *orb* and *plp/+, orb*). To quantify the relative localization of PLP at centrosomes, we used custom Python code to batch calculate the total intensity of PLP signals within 2 μm of the centrosome across all genotypes after validating this approach upon comparison with manual quantification (**Figure S5**). Quantification of PLP recruitment to centrosomes shows ∼20% reduction in *orb* mutants, which although not significantly different by one-way ANOVA across all genotypes, is significantly different from the WT control by an unpaired t-test (**Figure 4B**, *p*<0.01). Consistent with a requirement for *orb* in PLP localization, we noted a more drastic reduction (∼40% less than WT) in the recombinant *plp*/+, *orb* mutants (**Figure 4A, B**). We reasoned that the apparent reduction in PLP recruitment to centrosomes might be attributed to the reduction in total PLP levels, suggesting increasing *plp* dosage should rescue PLP localization in *orb* embryos. To test this hypothesis, we expressed a functional *PLP-GFP* transgene capable of rescuing *plp* mutant phenotypes in the *orb* mutant background (Galletta et al., 2014; Lerit et al., 2015). PLP distribution is restored at *PLP-GFP*; *orb* centrosomes, suggesting PLP localization to centrosomes functions downstream of *orb* activity (**Figure 4A, B**).

**Figure 4.**
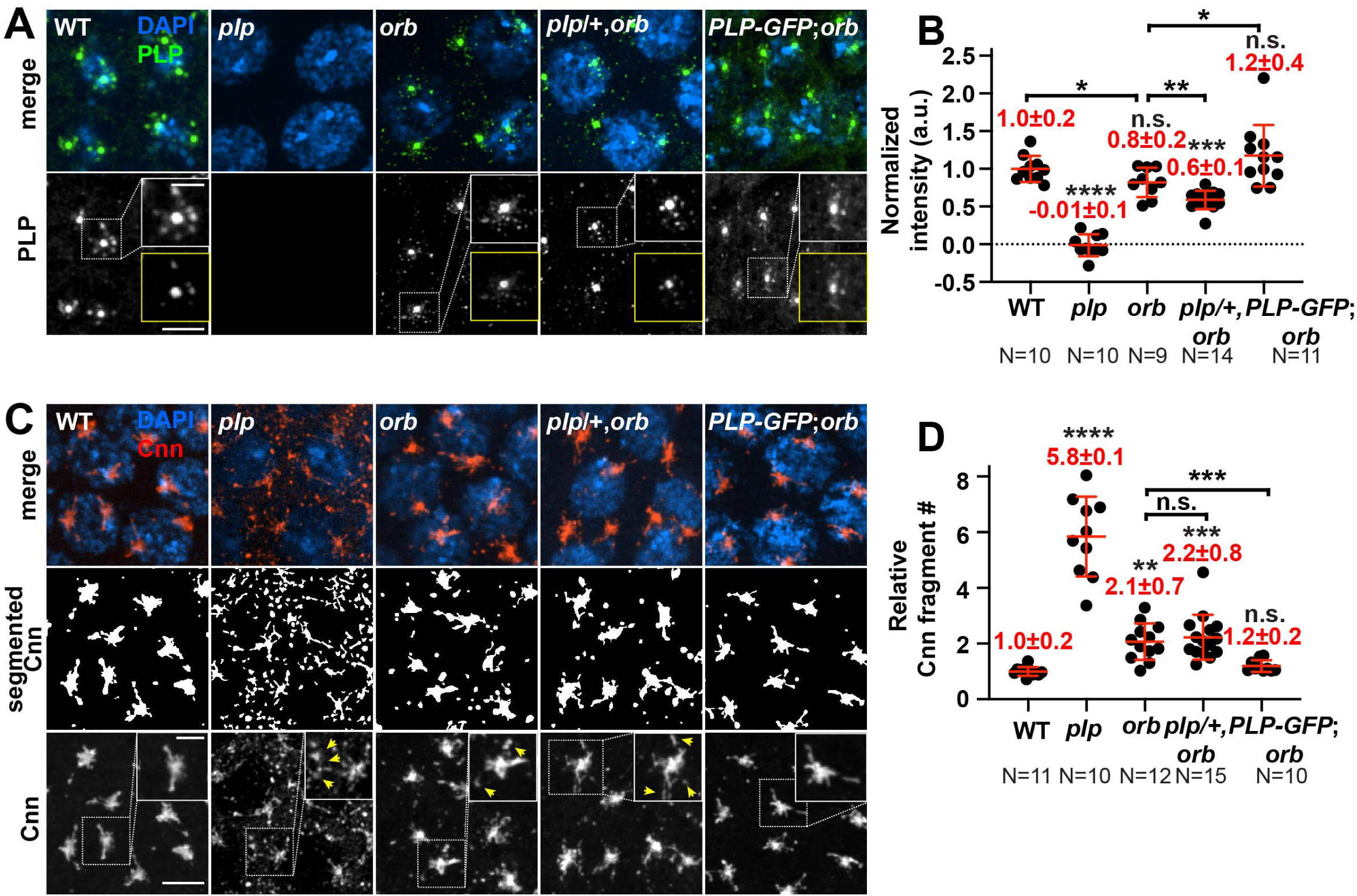
Formation of the PLP-Cnn PCM scaffold requires Orb. (A) Maximum intensity projections of NC 13 embryos of the indicated genotypes stained with anti-PLP antibodies (green). The LUT was adjusted to highlight PLP within the flare-zone. Boxed regions are enlarged in the inset, and less saturated images are shown in the yellow boxes. (B) Quantification of PLP intensity within 2 μm of the centrosome. Values were normalized to the mean intensity from WT embryos. Each dot represents the PLP intensity of one embryo averaged from all centrosomes in a 65 μm^2^ region. (C, top and bottom) Images show maximum intensity projections of NC 13 embryos stained with anti-Cnn antibodies (red). Yellow arrowheads highlight Cnn fragments. (C, middle row) Segmented Cnn images. (D) Quantification of the number of Cnn fragments. Values were normalized to the mean from WT embryos. Each dot represents the relative number of Cnn fragments of one embryo averaged from all centrosomes in a 65 μm^2^ region. Significance was determined by one-way ANOVA followed by Dunnett’s T3 multiple comparisons test relative to WT; bracket marks unpaired t-test. *orb* mutant: *orb^F343^*/*orb^mel^*; *plp*/+,*orb* mutant: *plp^2172^*/+, *orb^F343^*/*orb^mel^* ; *PLP-GFP*;*orb* mutant: *PLP-GFP*; *orb^F343^*/*orb^mel^*. Mean ± S.D. are displayed. n.s, not significant; \**p*<0.05; \*\**p*<0.01; \*\*\**p*<0.001; \*\*\*\**p*<0.0001. Bars: 5 μm; 2 μm (insets).

Our prior work showed PLP interacts directly with and organizes Cnn at centrosomes to scaffold the PCM (Lerit et al 2015). Given our observation that PLP levels and localization were diminished in *orb* mutants, we next addressed whether Cnn organization was affected, too. In control embryos, Cnn radiates symmetrically from centrosomes, forming interphase-specific flares, as previously described (**Figure 4C,** WT)(Megraw et al., 2002). Consistent with our prior work, Cnn distribution appears severely disorganized in *plp* embryos with fragmented Cnn flares and Cnn puncta dispersed within the cytosol (**Figure 4C,** *plp*) (Lerit et al., 2015; Fang and Lerit, 2020). Quantification of the relative number of cytoplasmic Cnn puncta in segmented images shows a significant, nearly 6-fold increase in Cnn puncta in *plp* embryos, as compared to WT (**Figure 4D**). Similar responses, albeit with reduced magnitude, were also observed in *orb* and *plp*/+, *orb* mutant embryos (**Figure 4C, D**). Moreover, expression of the *PLP-GFP* transgene rescues Cnn organization in *orb* mutants (**Figure 4C, D**, *PLP-GFP; orb*). These data suggest that Orb contributes to proper PLP-Cnn scaffold organization by regulating PLP protein expression and localization.

### Orb-dependent regulation of PLP supports genome stability

A requirement for Orb in modulating PLP levels, localization, and activity predicts that loss of *orb* would compromise centrosome function in maintaining mitotic fidelity. To investigate whether *orb* depletion is associated with chromosome instability (CIN), we stained WT and *orb* mutant embryos with DAPI, the mitotic marker pH3, and Asl to label centrioles (Varmark et al., 2007). Upon examination of anaphase embryos, the chromosomes in WT samples completely separated and lagging chromosomes were infrequent (N=3/20 WT embryos with CIN; **Figure 5A, B**; WT). In contrast, elevated rates of lagging chromosomes or chromosomal bridges were observed in *orb* mutants (55%, N=11/20 *orb* embryos with CIN), represented by lagging chromosomes at the spindle equator with persistent pH3-staining in anaphase/telophase-stage embryos (**Figure 5A,B**; *orb*) (Fox et al., 2010; Vitre and Cleveland, 2012). Moreover, *PLP-GFP* expression restored the mitotic fidelity of *orb* mutant embryos, evident by decreased CIN rates (21%, N=5/24 *PLP-GFP; orb* embryos with CIN; **Figure 5A, B**). These data implicate a requirement for *orb* in preserving genome stability in a mechanism dependent upon PLP dosage.

**Figure 5.**
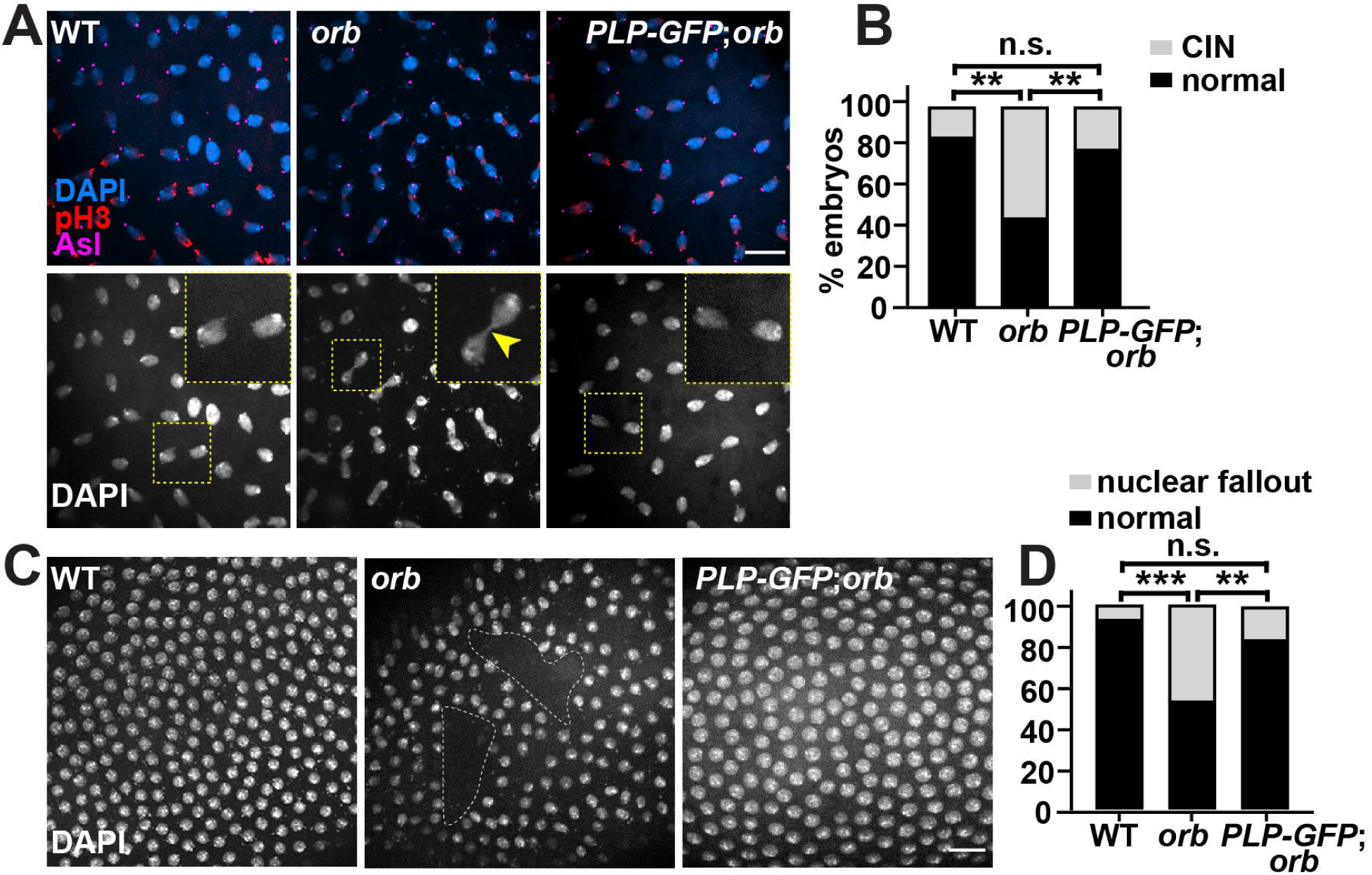
Orb functions upstream of PLP to support mitotic fidelity. (A) Images show maximum intensity projections of NC 12 telophase embryos labeled with anti-pH3 (red) and anti-Asterless (Asl, magenta) antibodies to visualize chromosome separation, as quantified in (B), where CIN indicates embryos with lagging chromosomes. (C) Maximum intensity projections of NC 13 interphase embryos labeled with DAPI to reveal NUF, as quantified in (D). \*\**p*<0.01; \*\*\**p*<0.001 by chi-square analysis. *orb* mutant: *orb^F343^*/*orb^mel^*; *PLP-GFP*;*orb* mutant: *PLP-GFP*; *orb^F343^*/*orb^mel^*. Bars: 20 μm.

To further examine genome stability, we measured rates of nuclear fallout (NUF), a developmental response to DNA damage in which damaged nuclei are ejected from the syncytial blastoderm cortex and targeted for apoptosis (Sullivan et al., 1993; Rothwell et al., 1998). NUF is readily detected as gaps in the normally uniform monolayer of DAPI-positive nuclei lining the embryonic cortex (dashed lines; **Figure 5C**). Although low rates of NUF are apparent even in WT embryos (7%, N=2/30), we observed significantly more NUF in *orb* mutants (47%, N=17/36; **Figure 5C, D**). However, NUF was largely rescued in *orb* mutants expressing the *PLP-GFP* transgene (16%, N=6/36 *PLP-GFP; orb* embryos with NUF; **Figure 5C, D**), further supporting the idea that Orb promotes genome stability via PLP.

Finally, to examine the mitotic fidelity of *orb* mutants in greater detail, we examined mitotic progression in live, cycling embryos expressing *GFP-γTub* to label the centrosomes and *H2A-RFP* to follow the chromosomes. As we were unable to recover double-balanced *orb^F343^* mutants, for these experiments we employed the weaker transheterozygous combination *orb^F303^*/*orb^mel^* to deplete Orb in our live imaging experiments (Lantz et al., 1994) (**Figure S2D**). While nuclei synchronously divided without NUF in most WT embryos (18%, N=2/11 showed NUF, **Figure 6A, Supplemental Video 1**), about 50% of *orb^F303^/orb^mel^* embryos displayed NUF (N=5/10 showed NUF, **Figure 6B, Supplemental Video 2**). Consistent with our quantification using fixed samples, expression of the *PLP-GFP* transgene largely ameliorated NUF in *orb* mutant embryos (17%, N=1/6 showed NUF, **Figure 6C, Supplemental Video 3**). Taken together, our results support a model wherein Orb contributes to centrosome function through regulating PLP expression.

**Figure 6.**
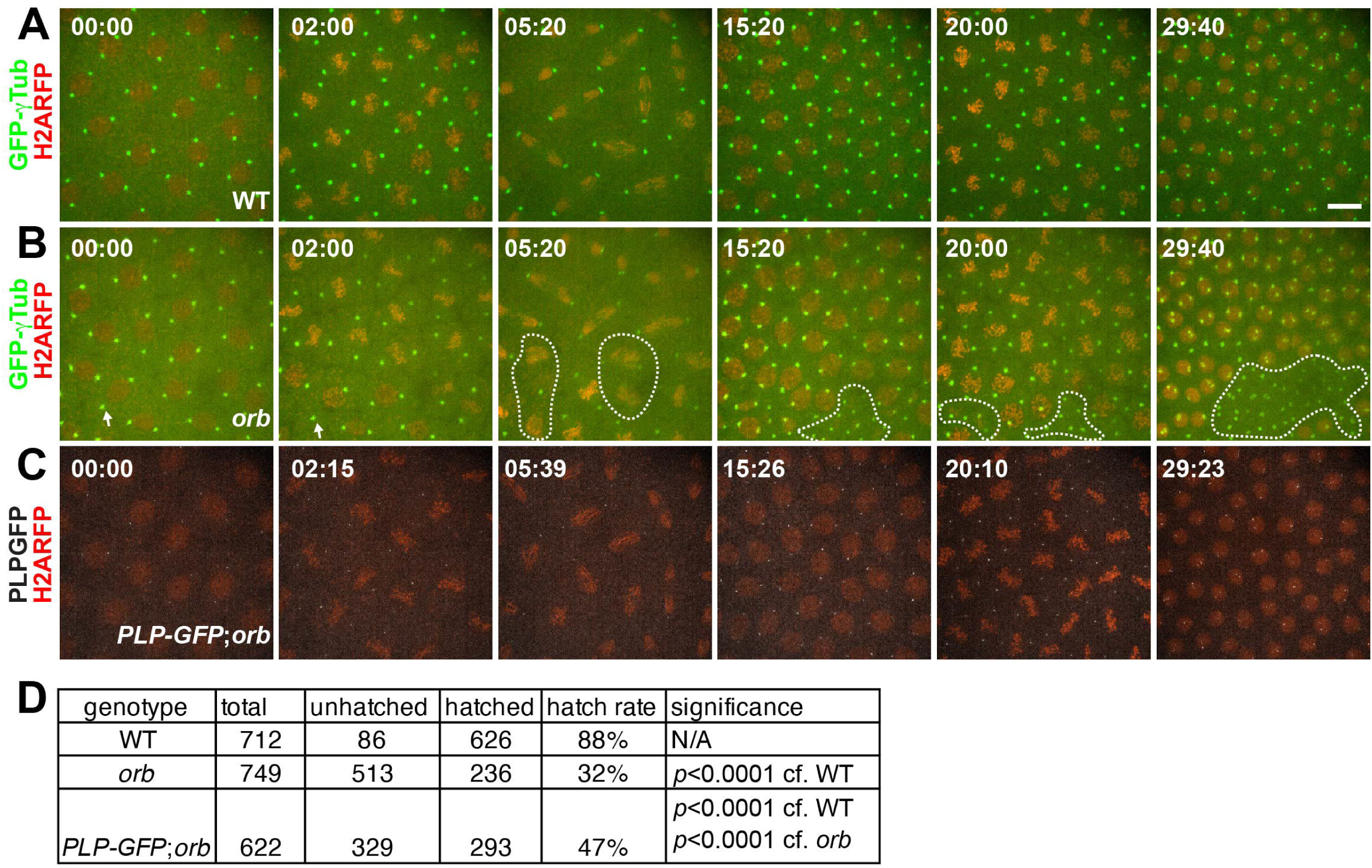
Orb promotes genome stability via PLP. Maximum intensity projected stills from time-lapse imaging of GFP-γTub or PLP-GFP with H2A-RFP in (A) control, (B) *orb^F303^*/*orb^mel^* mutant, and (C) *PLP-GFP*; *orb^F303^*/*orb^mel^* embryos. Time is in minutes. Dashed lines mark NUF. (D) Quantification of embryonic lethality rates from the indicated genotypes. Significance was determined by Fisher’s exact test. Bar: 10 μm. Supplemental Videos 1-3 are matched to panel (A-C) correspondingly.

Disruption of centrosome function impairs genome stability, which typically manifests in elevated rates of embryonic lethality. We examined embryonic viability in *orb* mutants and controls through hatch-rate analysis. Relatively low rates of embryonic lethality were noted in WT embryos, as nearly 90% hatched to first-instar larvae. In contrast to WT and consistent with previous studies, we observed significant rates of embryonic lethality in *orb^F343^/orb^mel^* mutants, as only 32% hatched to first-instar larvae (Christerson and McKearin, 1994; Lantz et al., 1994). Embryonic lethality in *orb* mutants is largely attributed to the requirement of Orb in establishing the embryonic patterning axes via its role in mediating localization and translation of *osk* and *grk* mRNAs (Christerson and McKearin, 1994; Chang et al., 1999; Chang et al., 2001; Castagnetti and Ephrussi, 2003). Interestingly, expression of the *PLP-GFP* transgene partially restored embryonic viability (53% hatched, *p*<0.0001, *orb* vs. *PLP-GFP*; *orb* by chi-squared test), supporting a genetic interaction between *orb* and *plp* required for viability (**Figure 6D**). Orb likely further supports embryonic viability through regulating other mRNA targets or protein partners (Chang et al., 2001; Mansfield et al., 2002; Norvell et al., 2015). Nonetheless, our data argue that *plp* is a critical downstream target subject to Orb regulation.

### Orb down-regulates *plp* RNA expression throughout oogenesis to embryogenesis

To investigate whether Orb regulates *plp* mRNA expression to promote protein expression, we performed qRT-PCR using primers designed to recognize all predicted *plp* mRNA variants. We detected a 1.3-fold increase in *plp* mRNA in 0-2 hr *orb* depleted embryos relative to WT (**Figure S6A**). Thus, the reduced levels of PLP protein observed in *orb* mutants may not be attributed to reduced *plp* mRNA transcription or RNA stability. During the first two hours of embryogenesis, zygotic transcription is largely inhibited, and RNA is mainly acquired from maternal oocytes (Tadros and Lipshitz, 2009). To test the hypothesis that the increased *plp* mRNA levels observed in *orb* embryos might be inherited from oogenesis, we examined *plp* mRNA in 2–4-day ovaries. *orb* ovarian extracts contain ∼1.4-fold more *plp* mRNA than WT, suggesting Orb normally attenuates *plp* mRNA expression or stability and consistent with the idea that the elevated levels of *plp* mRNA in *orb* mutants are passaged to the early embryo (**Figure S6B**). In contrast, we found PLP protein levels are unaltered in *orb* mutant ovarian extracts as compared to WT, indicating Orb-dependent regulation of PLP translation is apparently restricted to embryogenesis (**Figure S6C-D**).

### Orb facilitates the polyadenylation of *plp* mRNA

CPEB proteins, including Orb, mediate translational activation and/or repression by regulating the cytoplasmic polyadenylation of target mRNAs (Chang et al., 1999; Chang et al., 2001; Castagnetti and Ephrussi, 2003; Kim and Richter, 2007; Novoa et al., 2010; Burns et al., 2011; Norvell et al., 2015; Rojas-Ríos et al., 2015). To examine if Orb regulates *plp* mRNA polyadenylation to promote PLP translation, we monitored *plp* poly(A) tail length. Among the twelve *plp* mRNA variants, only *plp^RM^* uses the long 3’UTR (591 nt). The remaining 11 *plp* mRNA variants use a short 3’UTR (431 nt) (**Figure S2B; Fig 7A)** (Graveley et al., 2011). We first assayed which *plp* 3’UTR forms are expressed in 0-2 hr embryos by amplifying *plp* mRNA using one of two reverse primers: one aligns within a region common to both *plp* 3’UTRs (*plp*_Rev1_), while the other specifically aligns to the long *plp* 3’UTR (*plp*_Rev2_) (**Figure 7A**). PCR products corresponding to the *plp* 3’UTR were further validated by restriction enzyme digestion. While both *plp* 3’UTRs are sensitive to BmrI digestion, only the long *plp* 3-UTR is sensitive to EcoR1 (Fig. 7A, B, lanes 1–8). Although our PCR profiling confirms expression of the long 3’UTR associated with *plp^RM^* in 0-2 hr embryos, bioinformatic analysis of published RNA-seq datasets, indicates *plp^RM^* has a lower expression level than other *plp* variants in early embryos (Graveley et al., 2011).

**Figure 7.**
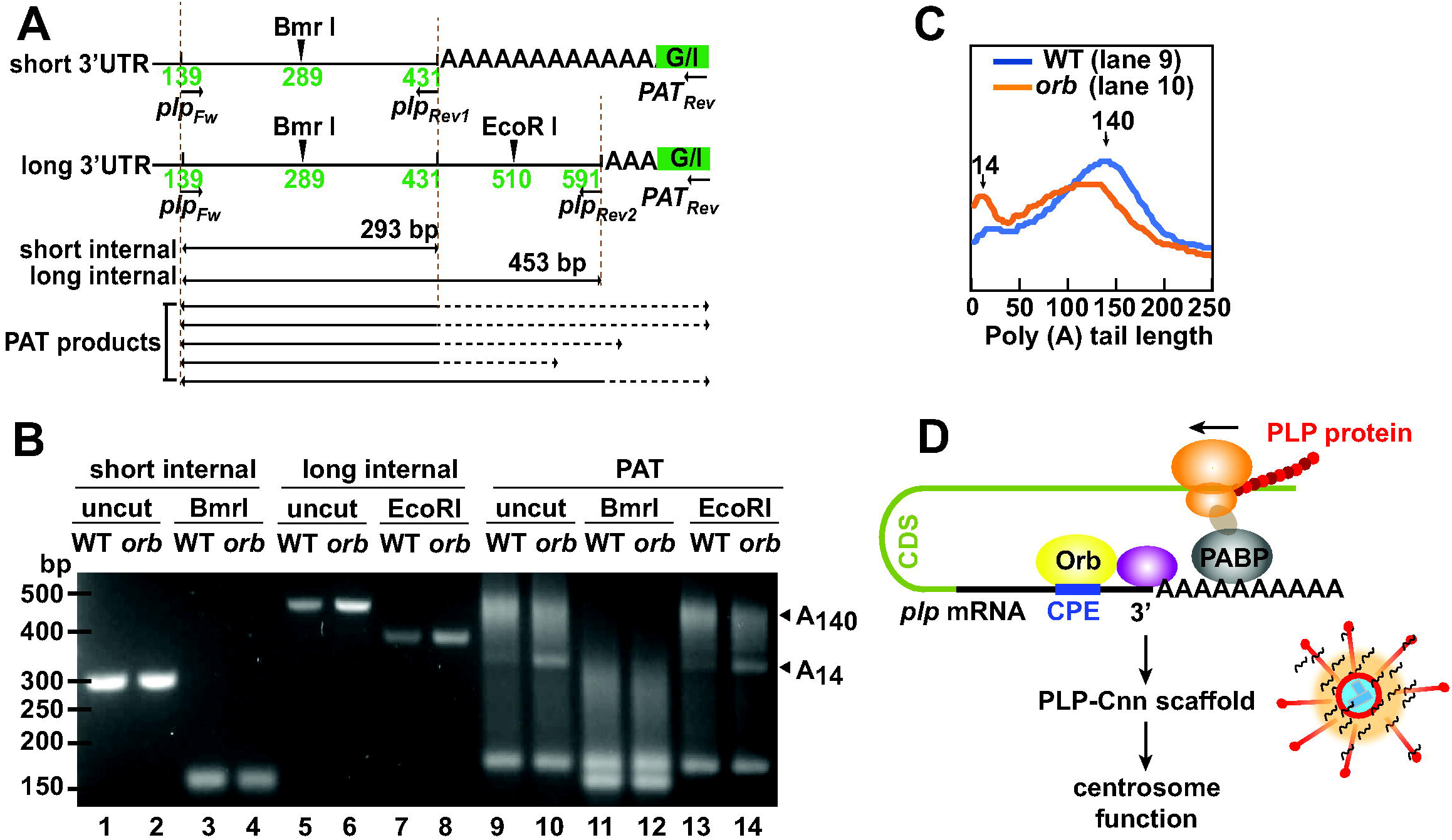
Orb promotes *plp* polyadenylation. (A) Schematic shows positions of PCR primers and the EcoRI and BmrI restriction sites on the short and long *plp* 3’UTRs. The predicted sizes of internal PCR products are listed below. (B) Gel shows cDNA products amplified from G/I- tailed RNA extracted from 0–2-hr control (*iso-1*) and *orb^F343^*/*orb^mel^* embryos. PCR products were digested with EcoRI and/or BmrI, as indicated. The approximate lengths of poly(A)-tails are indicated (arrowheads). (C) Line profiles from lanes 9 and 10. Poly(A)-tail length was calculated by subtracting the internal PCR product plus G/I tail length from the total PAT product length. (D) A model for Orb-mediated regulation of *plp* mRNA through promoting *plp* polyadenylation. Orb interacts with *plp* mRNA, possibly through CPE motifs in the 3’UTR. Orb may further recruit poly(A) polymerase (purple) to the *plp* 3’UTR to enable polyadenylation. The elongated poly(A)- tail likely stabilizes the interaction between *plp* mRNA and PABP, which can further recruit additional factors (brown) required for ribosome (orange) assembly and PLP protein synthesis. PLP expression supports PLP-Cnn scaffold assembly necessary for centrosome function and error-free mitosis. Uncropped gel is available to view on Figshare: https://doi.org/10.6084/m9.figshare.16900471.v1

We next performed a poly(A) test (PAT) assay of *plp* mRNA from control versus *orb^F343^/orb^mel^* mutant embryos (Legnini et al., 2019). For this, we tagged polyadenylated mRNA from 0-2 hr embryos with guanosine and inosine (G/I-tailing), followed by RT-PCR to synthesize cDNA. *plp* poly(A) tails were then amplified by PCR using a universal reverse primer and a *plp*- specific forward primer (**Figure 7A**). PAT products resolved as a ∼339–525 bp smear by gel electrophoresis (**Figure 7B,** lanes 9–10), corresponding to approximately 14-200 nt poly(A) length (**Figure 7B**, arrowheads). We used restriction enzyme digestion to distinguish PAT products from the *plp* long versus short 3’-UTRs. As *plp* PAT products are sensitive to BmrI digestion, but only modestly altered by EcoRI, we conclude polyadenylated products from the short *plp* 3’UTR are more enriched in 0-2 hr embryos than the long *plp* 3’-UTR (**Figure 7B, lane 9–14**). Plotting a line profile of the PAT products detected in control versus *orb* embryos reveals a leftward shift indicative of reduced polyadenylation of *plp* mRNA in *orb* mutants (**Figure 7C**). Because polyadenylation stimulates translation (Eichhorn et al., 2016; Passmore and Coller, 2021; Xiang and Bartel, 2021), our results are consistent with a model wherein Orb activates *plp* mRNA translation by promoting polyadenylation. The elongated poly(A) tail may stabilize the interaction between *plp* mRNA and poly(A)-binding protein (PABP), which can further recruit proteins required for ribosome assembly and PLP protein synthesis (**Figure 7D**).

## Discussion

We recently analyzed the distributions of five centrosomal mRNAs, including *plp* mRNA, revealing a common cell cycle-dependent enrichment at late interphase centrosomes (Ryder et al., 2020). Our present work extended this analysis, highlighting enrichment of *plp* mRNA at centrosomes across syncytial development. One simple model is that centrosomal mRNAs are more likely to localize to interphase centrosomes, which are larger than mitotic centrosomes in syncytial *Drosophila* embryos (Megraw et al., 2002; Lerit et al., 2015). This model oversimplifies the specificity of RNA localization to centrosomes – relatively few RNAs reside at centrosomes (Lécuyer et al., 2007; Chouaib et al., 2020; Kwon et al., 2021; Safieddine et al., 2021). Moreover, similar interphase/prophase-stage preferential enrichments of centrosomal mRNAs were recently observed in cultured human cells, wherein centrosome size is larger in mitosis (Sepulveda et al., 2018; Safieddine et al., 2021). Collectively, these studies demonstrate that multiple mRNAs across divergent species co-enrich at centrosomes at the same cell cycle phase, suggestive of conserved localization mechanisms. How RNA localization to centrosomes is molecularly executed in a cell cycle-dependent manner is a fascinating question requiring further investigation.

Toward this end, we identified conservation of multiple CPE sites within the *plp* and *PCNT* 3’UTRs. Because CPE-containing mRNAs are enriched at spindle poles and CPEB is required for centrosomal localization of *Cyclin B1* mRNA localization, we reasoned CPE sites may contribute to *plp*, and possibly *PCNT*, mRNA localization. Given its established roles in mediating localization of other mRNAs, such as *osk* and *grk*, we hypothesized Orb might contribute to the enrichment of *plp* mRNA to centrosomes. Our data do not support this hypothesis. Strong hypomorphic *orb* mutant embryos did not show altered *plp* mRNA localization to centrosomes.

In cell culture models, *PCNT* mRNA localization to the centrosome is puromycin-sensitive, suggesting it is trafficked by a co-translational transport mechanism (Sepulveda et al., 2018; Safieddine et al., 2021). While *orb* depletion does not impair *plp* mRNA localization to centrosomes, it does result in a 50% reduction in total PLP levels, suggesting RNA localization and translation may be uncoupled. It is feasible that multiple *orb*-independent mechanisms may contribute to *plp* mRNA localization to centrosomes.

Our data support a model whereby Orb regulates PLP translational activation by facilitating polyadenylation. In many organisms, the first hours of embryogenesis rely upon maternally provided mRNAs and proteins until the maternal-to-zygotic transition activates bulk zygotic transcription (Tadros and Lipshitz, 2009). Consequently, translational regulation is a signature paradigm of gene regulation during early development (Johnstone and Lasko, 2001). Because syncytial *Drosophila* embryos divide every 10-20 minutes, they undergo rapid, successive centrosome doubling events and oscillations in PCM recruitment and shedding at compressed timescales. Here, we define a mechanism of translational activation of a conserved PCM scaffolding factor, PLP, mediated by the CPEB member, Orb.

We detected differential usage among two alternative 3’UTRs, showing the shorter *plp* 3’UTR is favored during embryogenesis. Moreover, monitoring poly(A)-tail length revealed reduced polyadenylation at the shorter *plp* 3’UTR in *orb* mutants as compared to controls. Genome-wide studies reveal mRNA poly(A) length is generally positively correlated to translation efficiency in *Drosophila* mature oocytes and early embryos (Subtelny et al., 2014; Eichhorn et al., 2016; Lim et al., 2016). Other studies using luciferase reporter systems also demonstrated that poly(A)-tail length strongly influences translation efficiency (Coll et al., 2010; Xiang and Bartel, 2021). Taken together, our data suggest polyadenylation by Orb normally promotes *plp* mRNA translation. Consistent with this model, the poly(A)-tail length of the Orb target mRNAs *osk* and *Atg12* correlate with translational efficiency (Castagnetti and Ephrussi, 2003; Rojas-Ríos et al., 2015).

The mechanism by which CPEB proteins promote translation is best defined in *Xenopus* oocytes, where phosphorylated CPEB1 recruits the cleavage and polyadenylation specificity factor (CPSF) and poly(A) polymerase (PAP) to the 3’UTR of mRNA targets and elongates their poly(A)-tails. The polyadenylated mRNA then recruits and stabilizes interactions with poly(A)- binding protein (PABP) and further recruits eukaryotic translation initiation factors (EIFs), leading to ribosome assembly and protein translation (Mendez et al., 2000b).

While PLP protein levels are diminished in *orb* mutants, PLP localization to centrosomes is more modestly impaired. We detected ∼20% less PLP localizing to interphase centrosomes in NC 13 *orb* embryos versus ∼50% less PLP protein in embryonic extracts pooled across NC 1– NC 14. This variance may be attributed to cell cycle or NC-specific dynamics of PLP protein expression or localization. In addition, perdurance of PLP protein at the centrosome may be due to its relatively slow rate of turnover (Conduit et al., 2014; Richens et al., 2015). Our data suggest that while Orb supports PLP protein localization to centrosomes, redundant mechanisms exist to ensure robust localization, consistent with the integral role PLP plays in PCM scaffolding.

## Materials and methods

### Fly stocks

The following stocks and transgenic lines were used: *y^1^w^1118^* (Bloomington Drosophila Stock Center (BDSC) #1495) was used as the WT control unless otherwise noted. The genome reference genotype *iso-1* (BDSC #2057) was used as the control for PAT assay (Brizuela et al., 1994). In all experiments except for live imaging, *orb* mutant embryos were *orb^F343/^orb^mel^* transheterozygotes (*orb^F343^*; BDSC #58477; (Lantz et al., 1994) and *orb^mel^*; BDSC #58743; (Christerson and McKearin, 1994)). For live imaging, *orb^F303/^orb^mel^* transheterozygotes were used (Christerson and McKearin, 1994; Lantz et al., 1994). Null *Fmr1* mutant embryos were *Fmr1^△113M^/Fmr1^3^* transheterozygotes (*Fmr1^△113M^*; BDSC #67403 and *Fmr1^3^*, gift from T.Jongens, University of Pennsylvania; (Dockendorff et al., 2002)). Null *orb2* mutant embryos were *orb2^36^/orb2^7^* transheterozygotes (*orb2^7^*; BDSC #58480 and *orb2^36^*; BDSC #58479; (Xu et al., 2012)); *plp^2172^* allele is a null allele (Spradling et al., 1999; Martinez-Campos et al., 2004) and was recombined onto the *orb^F343^* chromosome to generate *plp^2172^, orb^F343^* recombinant animals. *Ubi-GFP-γ-Tub23C* expresses GFP-γTub under the *Ubiquitin* (*Ubi*) promotor (Lerit and Rusan, 2013); *Ubi-PLP^FL^-GFP* expresses full-length PLP isoform PF under the *Ubi* promoter (Galletta et al., 2014); *H2A-RFP* expresses a red fluorescent H2A variant under endogenous regulatory elements (Pandey et al., 2005); and *GFP-Orb* is a gene trap expressing a GFSTF (EGFP, FlAsH, StrepII, TEV, and 3xFLAG) tag under endogenous regulatory elements (BDSC #59817; (Nagarkar-Jaiswal et al., 2015)). To examine maternal effects, mutant embryos are progeny derived from mutant mothers. Flies were raised on Bloomington formula ‘Fly Food B’ (LabExpress; Ann Arbor, MI), and crosses were maintained at 25°C in a light and temperature-controlled incubator chamber.

### Bioinformatics

To compare overlapping mRNA targets of Orb (Stepien et al., 2016) and CPEB1 (Pascual et al., 2021), *Drosophila* gene names were converted to human gene identifiers using the of Query symbols/IDs tool in Flybase (Larkin et al., 2021). Overlapping genes were identified by the ‘COUNT IF’ function in Excel. Venn diagrams were plotted using the Meta-Chart online tool (https://www.meta-chart.com/). GO cellular component analysis of common genes was analyzed using the Panther statistical overrepresentation test (http://www.pantherdb.org/), and Fisher’s exact test was used to generate an adjusted p-value, i.e., false discovery rate (FDR) (Mi et al., 2021).

To align human *PCNT* (NCBI; NM_006031.6) and *Drosophila plp* 3’UTRs, Clustal Omega was used (Madeira et al., 2019). Conservation of CPE motifs with the *plp* 3’UTRs from different *Drosophila* species was compared using the conservation 124 insect track on the UCSC genome browser (Kent et al., 2002).

### Embryonic hatch rate analysis

6–12 hr eggs were collected on yeasted grape juice agar plates and ∼200 embryos were transferred to fresh plates and aged for 48 hr at 25°C. Unhatched embryos were counted from each plate. Three independent replicates were performed.

### Live imaging

Embryos were prepared for live imaging as described in (Lerit et al., 2017). Briefly, dechorionated embryos were adhered to a sticky 22 × 30 mm # 1.5 glass coverslip, covered with a thin layer of halocarbon oil, and inverted onto a clear, gas-permeable 50 mm dish using broken #1 glass coverslips as spacers. Images were captured at 1 μm z-intervals over a 10–15 μm volume at 20 s intervals.

### Immunofluorescence

For immunofluorescence with PLP and Cnn antibodies, embryos were fixed in a 1:1 solution of anhydrous methanol (Sigma, #322415): heptane for 15 sec and devitellinized in methanol. For smFISH and IF staining with Asl and pH3 antibodies, embryos were fixed in a 1:4 solution of 4% paraformaldehyde: heptane for 20 min and devitellinized in methanol (Rothwell and Sullivan, 2007). Fixed embryos were rehydrated, blocked in BBT buffer (PBS supplemented with 0.1% Tween-20 and 0.1% BSA), and incubated overnight at 4°C with primary antibodies diluted in BBT. For Asl staining, BBT buffer with 0.5% BSA was used. After washing, embryos were further blocked in BBT supplemented with 2% normal goat serum and incubated for 2 hr at room temperature with secondary antibodies and DAPI (10 ng/ml, ThermoFisher). Embryos were mounted in Aqua-Poly/Mount (Polysciences, Inc.) prior to imaging.

The following primary antibodies were used: rabbit anti-PLP antibody (1:4000, gift from N. Rusan, NIH), guinea pig anti-Asl (1:4000, gift from G. Rogers, University of Arizona), rabbit anti-Cnn (1:4000, gift from T. Megraw, Florida State University), mouse anti-phospho-Histone H3 Ser10 (pH3; 1:1000, Millipore 05570). Secondary antibodies: Alexa Fluor 488, 568, or 647 (1:500, Molecular Probes).

### smFISH detection and analysis

Stellaris *plp* and *gapdh* smFISH probes conjugated to Quasar 570 dye (LGC Biosearch Technologies; Middlesex, UK) were designed against the coding region for each gene using the Stellaris RNA FISH probe designer (Ryder et al., 2020; Ryder and Lerit, 2020). smFISH probes were dissolved in nuclease-free water at 25 μM and stored at -20°C before use.

smFISH experiments were performed as previously described (Ryder et al., 2020; Ryder and Lerit, 2020). All the following steps were performed with RNase-free solutions. Embryos were rehydrated and washed first in 0.1% PBST (PBS plus 0.1% Tween-20) and then in wash buffer (WB; 10% formamide and 2× SSC supplemented fresh each experiment with 0.1% Tween-20 and 2 μg/ml nuclease-free BSA). Embryos were then incubated with 100 μl of hybridization buffer (HB; 100 mg/ml dextran sulfate and 10% formamide in 2× SSC supplemented fresh each experiment with 0.1% Tween-20, 2 μg/ml nuclease-free BSA, and 10 mM ribonucleoside vanadyl complex (RVC; S1402S; New England Biolabs; Ipswich, MA) for 10–20 min in a 37°C water bath. Embryos were then incubated in 25 μl of HB containing 0.5 μM smFISH probes in a 37°C water bath overnight. Embryos were washed three times for 30 min in prewarmed WB, stained with DAPI (1:1000) for 1 hr at room temperature, washed with 0.1% PBST, and mounted with Vectashield mounting medium (H-1000; Vector Laboratories; Burlingame, CA). Slides were stored at 4°C and imaged within 1 week.

smFISH signals were detected and single molecule normalization was performed as recently described (Ryder et al., 2020; Ryder and Lerit, 2020). Briefly, single-channel .tif raw images were segmented in three dimensions using Python scripts adapted from the Allen Institute for Cell Science Cell Segmenter (Chen et al., 2018). Each segmented image was compared with the raw image to validate accurate segmentation. RNA objects of ≥50 pixels in segmented images were identified, and object features were extracted, which included surface coordinates. Distances were measured from the surface of each RNA object to the surface of the closest centrosome. We calculated the percentage of total RNA within 1 μm from the centrosome surface and selected 10, 8, 6 and 4 μm as the upper boundary for the pseudo-cell radius for NC 10, NC 11, NC 12, and NC 13; respectively, based on measuring the centrosome-to-centrosome distances from a set of representative images. Later interphase/prophase embryos were selected by their large, round nuclei and metaphase samples were identified by alignment of condensed chromosomes at the metaphase plate.

### Microscopy

Images were acquired on a Nikon Ti-E system fitted with a Yokagawa CSU-X1 spinning disk head, Hamamatsu Orca Flash 4.0 v2 digital complementary metal oxide-semiconductor (CMOS) camera, Perfect Focus system (Nikon), Nikon LU-N4 solid state laser launch (15 mW 405, 488, 561, and 647 nm) using a Nikon 100x, 1.49 NA Apo TIRF oil immersion objective. The microscope was powered through Nikon Elements AR software on a 64-bit HP Z440 workstation.

### Image analysis

Fiji (National Institutes of Health; (Schindelin et al., 2012)) was used for all image analysis. To examine nuclear fallout, embryos were stained with anti-Asl antibody to label centrosomes and DAPI to label DNA (Lerit et al., 2015). Areas where centrosomes clustered but were devoid of nuclei were counted as sites of nuclear fallout from genotype-blinded images. To examine CIN, embryos were stained with anti-Asl to label centrosomes, anti-pH3 to label anaphase lagging chromosomes, and DAPI to label DNA. Anaphase-stage embryos were scored for lagging chromosomes from genotype-blinded images, as defined by laggard chromosome(s) at the spindle equator showing persistent pH3-staining (Fox et al., 2010).

To quantify PLP localization to centrosomes, single-channel PLP .tif raw images were processed in batch across all genotypes. Blinded images were segmented in three dimensions to display all PLP objects (as segmentation 1), including centriolar and flare zone signals, using Python scripts adapted from the Allen Institute for Cell Science Cell Segmenter (Chen et al., 2018). Each segmented image was compared with the original image to validate accurate segmentation. A second segmentation was performed to only display the PLP objects at centrioles (as segmentation 2), used as a reference point to identify all PLP objects (segmentation 1) within a centrosome (segmentation 2). Next, all PLP objects were identified, and the raw image total pixel intensity was extracted from a 2 μm region to calculate the intensity of all PLP objects at the centrosome. To validate the accuracy of batch-analysis for PLP quantification, we compared background-subtracted PLP intensity measurements (integrated densities) within 2 μm of the centrosome from N=10 maximum-intensity projected NC 13 WT and recombinant *plp/+,orb* embryos using manual quantification. Similar results were obtained (Fig. S5), and batch analysis was used subsequently (Fig. 4).

To quantify the number of Cnn fragments per pseudo-cell, single-channel Cnn .tif raw images were segmented as above. We then used the 3D Objects Counter tool in Fiji to quantify the total number of Cnn fragments in the segmented images, which was then averaged by the total number of nuclei.

Images were assembled using Fiji, Adobe Photoshop, and Adobe Illustrator software to separate or merge channels, crop regions of interest, generate maximum-intensity projections, and adjust brightness and contrast.

### RNA-immunoprecipitation

RNA-immunoprecipitation was performed as previously described in (Ryder et al., 2020). Briefly, ovaries were dissected from ∼20 well-fed females and homogenized on ice in 200 μl lysis buffer (50 mM Hepes, pH 7.4, 150 mM NaCl, 2.5 mM MgCl_2_, 250 mM sucrose, and 0.1% Triton X-100) supplemented with 1x EDTA-free protease inhibitor cocktail (Roche, 04693159001), 1 μg/mL Pepstatin A (Sigma, P5318), 1 mM DTT (Sigma, 10197777001), 1 U/μl RNase inhibitor (M0314S; New England Biolabs), and 2 mM RVC. Lysates were cleared by centrifugation at 12,000 rpm, and the supernatant was precleared in 20 μl binding control magnetic beads (bmp-20, ChromoTek; Munich, DE) for 30 min at 4°C. Precleared supernatant was then immunoprecipitated with GFP-Trap magnetic agarose beads (gtma-10, ChromoTek) for 2 hr at 4°C. Beads were then washed in lysis buffer supplemented with 0.4 mM RVC and resuspended in 100 μl lysis buffer. 25 μl of bead slurry was reserved and analyzed for protein content by western blotting. RNA was extracted from the remaining 75 μl of beads TRIzol Reagent (#15596026, ThermoFisher Scientific) following treatment with 1 μl TURBO DNase (#AM1907; ThermoFisher Scientific).

cDNA was synthesized from 500 ng of RNA using Superscript IV Reverse Transcriptase (18091050; Thermo Fisher Scientific) according to the manufacturer’s protocol. DNA was amplified by PCR using Phusion High Fidelity DNA Polymerase (M0530L; New England Biolabs) using the following primers:

*plp* Forward: GAAGCCATATCGAAGACACTC
*plp* Reverse: TGTCAGCCAATAGTCAGTCG
*gapdh* Forward: CACCCATTCGTCTGTGTTCG
*gapdh* Reverse: CAACAGTGATTCCCGACCAG
*orb* Forward: GTGTGAGACTTTGGACTTGTAGG
*orb* Reverse: GTTTCGATTCGAGGGTGTTCG

### Immunoblotting

Embryo extracts were prepared from ∼10 μl of methanol-fixed embryos. Embryos were rehydrated with 0.1% PBST and homogenized in 150 μl 0.1% PBST using a disposable plastic pestle and cordless motor. Ovarian extracts were similarly prepared from ovaries dissected from 10 well-fed females. 30 μl of 5X SDS loading buffer was added into the protein extracts, and the samples were boiled at 95°C for 10 min. Extracts were stored at -20°C or immediately resolved on an 7.5% SDS-PAGE pre-cast gel (Bio-Rad, #4568023) and transferred onto a 0.2 μm nitrocellulose membrane (GE healthcare, #10600001) by wet-transfer in a buffer containing 25 mM Tris-HCl, pH 7.6, 192 mM glycine, 10% methanol, and 0.02% SDS. Membranes were blocked for 1 hour at room temperature in 5% dry milk diluted in TBST (Tris-based saline with 0.05% Tween-20), washed well with TBST, and incubated overnight at 4 °C with primary antibodies. After washing with TBST, membranes were incubated for 1.5 hr in secondary antibodies diluted 1:5000 in TBST. Bands were visualized with Clarity ECL substrate (Bio-Rad, 1705061) on a Bio-Rad ChemiDoc imaging system.

The following primary antibodies were used: rabbit anti-PLP (1:4000, gift from N. Rusan, NIH), mouse anti-Khc SUK 4 (1:200, DSHB; J.M. Scholey, University of Colorado), mouse anti-β-Tubulin E7 (1:1000, DSHB; M. Klymkowsky, University of Colorado), mouse anti-Orb 4H8 (1:100, DHSB; P. Schedl, Princeton University). Secondary antibodies: goat anti-mouse HRP (1:5000, ThermoFisher #31430) and goat anti-rat IgG HRP (1:5000, Caltag Medsystems #6908-250). Densitometry was measured in Adobe Photoshop and protein levels are normalized to the loading control.

### qRT-PCR

RNA was extracted from ∼2-5 mg of dechorionated 0–2 hr embryos or ovaries dissected from 5 well-fed females per biological replicate using TRIzol Reagent (#15596026, ThermoFisher Scientific) and treated with1 μl TURBO DNase (2 U/μl, # AM1907, ThermoFisher Scientific) prior to RT-PCR. 500 ng of RNA was reverse transcribed to cDNA with Superscript IV Reverse Transcriptase following the manufacturer’s protocol. qPCR was performed on a Bio-Rad CFX96 Real-time system with iTaq Universal SYBR Green Supermix (#1725121, Bio-Rad; Hercules, CA). Values were normalized to *RpL32* (*rp49*) expression levels. Ct values from the qPCR results were analyzed and the relative expression levels for each condition were calculated using Microsoft Excel. Three biological replicates and three technical replicates were performed on a single 96-well plate.

The primers used in this study are listed below:

*rp49* Forward: CATACAGGCCCAAGATCGTG
*rp49* Reverse: ACAGCTTAGCATATCGATCCG
*plp* Foward: CGCAGCAAGGAGGAGATAAC
*plp* Reverse: TCAGCCTGCAGTTTGTTCAC

### Poly(A) tail test (PAT) assay

2 mg of 0-2 hr embryos were harvested and RNA was extracted with TRIzol Reagent (#15596026, ThermoFisher Scientific). The PAT test was conducted following the manufacturer’s protocol (ThermoFisher, PAT kit, #76455). Breifly, 3 μg of RNA was digested with Turbo DNase (2 U/μl, # AM1907, ThermoFisher Scientific). 500 ng of DNase-digested RNA was tagged with G/I tails in a 10 μl reaction, and 5 μl of G/I-tailed RNAs were reverse transcribed using a universal primer complementary to the G/I tails to synthesize cDNA in a 20 μl reaction. 20 μl of cDNA was diluted 1:1 into nuclease-free water, and 10 μl was then used to amplify PAT products using a *plp*-specific forward primer and a universal reverse primer complementary to the G/I tails in a 50 μl reaction using a two-step PCR protocol and Taq polymerase, as provided by the manufacturer. To amplify long or short *plp* 3’-UTRs, 2 μl of diluted cDNA was used as template using the indicated primers in a 50 μl reaction using Phusion High Fidelity DNA Polymerase (M0530L; New England Biolabs). 15 μl of amplified DNA was used for EcoRI and BmrI restriction enzyme digestion (New England Biolabs). DNA products were separated by gel electrophoresis on a 3% agarose gel. The primer PAT_Rev_ is from the manufacturer and is proprietary, the primers we designed that were used in this study are listed below:

*plp*_Rev1_: CGAATGTGAAATAAATTTGGTT
*pl*p_Rev2_: CTACTGCTTTCGATACCTTTTT
*plp*_Fw_: ACCTGTACCATTTCCCCTCA

### Statistical methods

Data were plotted and statistical analysis was performed using GraphPad Prism software. To calculate significance, distribution normality was first confirmed with a D’Agnostino and Pearson normality test. Data were then analyzed by Student’s two-tailed t-test or One-way ANOVA test and are displayed as mean ± SD or a chi-square test as indicated in the figure legends. Data shown are representative results from at least two independent experiments.

## Author contributions

DAL– conceptualization, funding acquisition, project administration, supervision, writing–original draft, and writing–review & editing.

JF– formal analysis, funding acquisition, investigation, methodology, project administration, software, validation, visualization, writing–original draft, and writing–review & editing.

## Acknowledgements

We thank Drs. Nasser Rusan, Greg Rogers, and Timothy Megraw for gifts of reagents; Drs. Elizabeth Gavis, Amanda Norvell, and Martine Simonelig for advice about the PAT assay; Dr. Pearl Ryder for computational assistance; Ms. Beverly Robinson for technical advice on ontological analysis; and Mr. Jovan Brockett for embryo collections. We are grateful to our colleagues for constructive feedback on the manuscript.

This work was supported by NIH grant R01GM138544 to D.A. Lerit and American Heart Association grant 20POST35210023 to J. Fang.

## Supplemental Data

**Figure S1.**
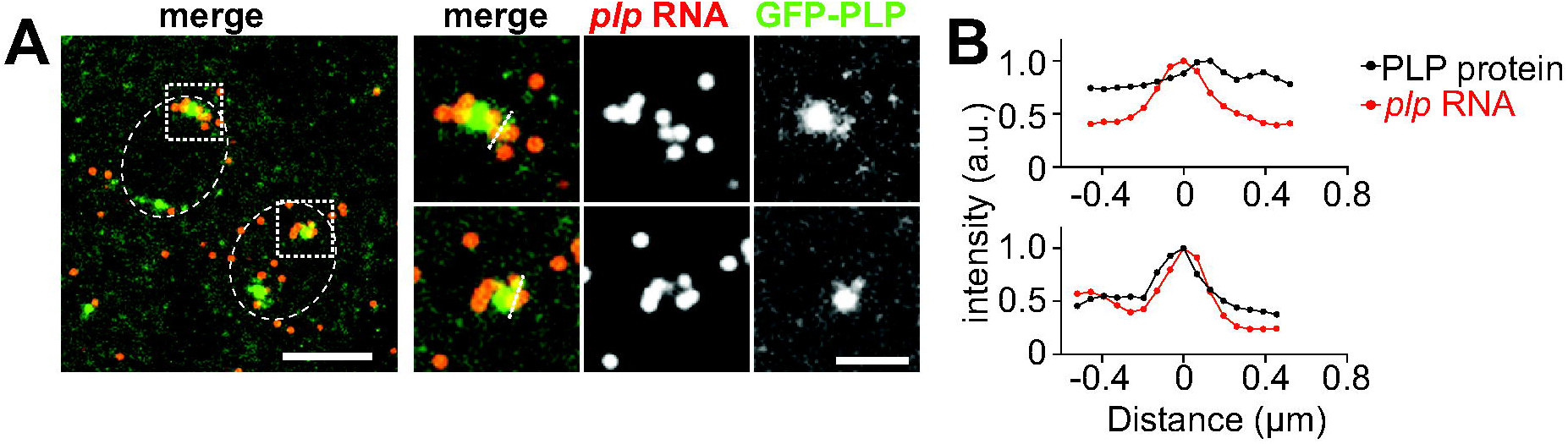
PLP protein overlaps with *plp* mRNA. (A) NC 13 *Drosophila* embryos expressing *GFP-PLP* (green) were stained with *plp* smFISH probes (red). Boxed regions are magnified in insets on the right. Dashed ovals mark the pseudo cells. (B) Plot profile analysis of GFP-PLP and *plp* mRNA from the centrosomes, as shown in insets of (A). Distances were normalized to the peak fluorescence intensity of *plp* mRNA. Bars: 5 μm; inset, 2 μm.

**Figure S2.**
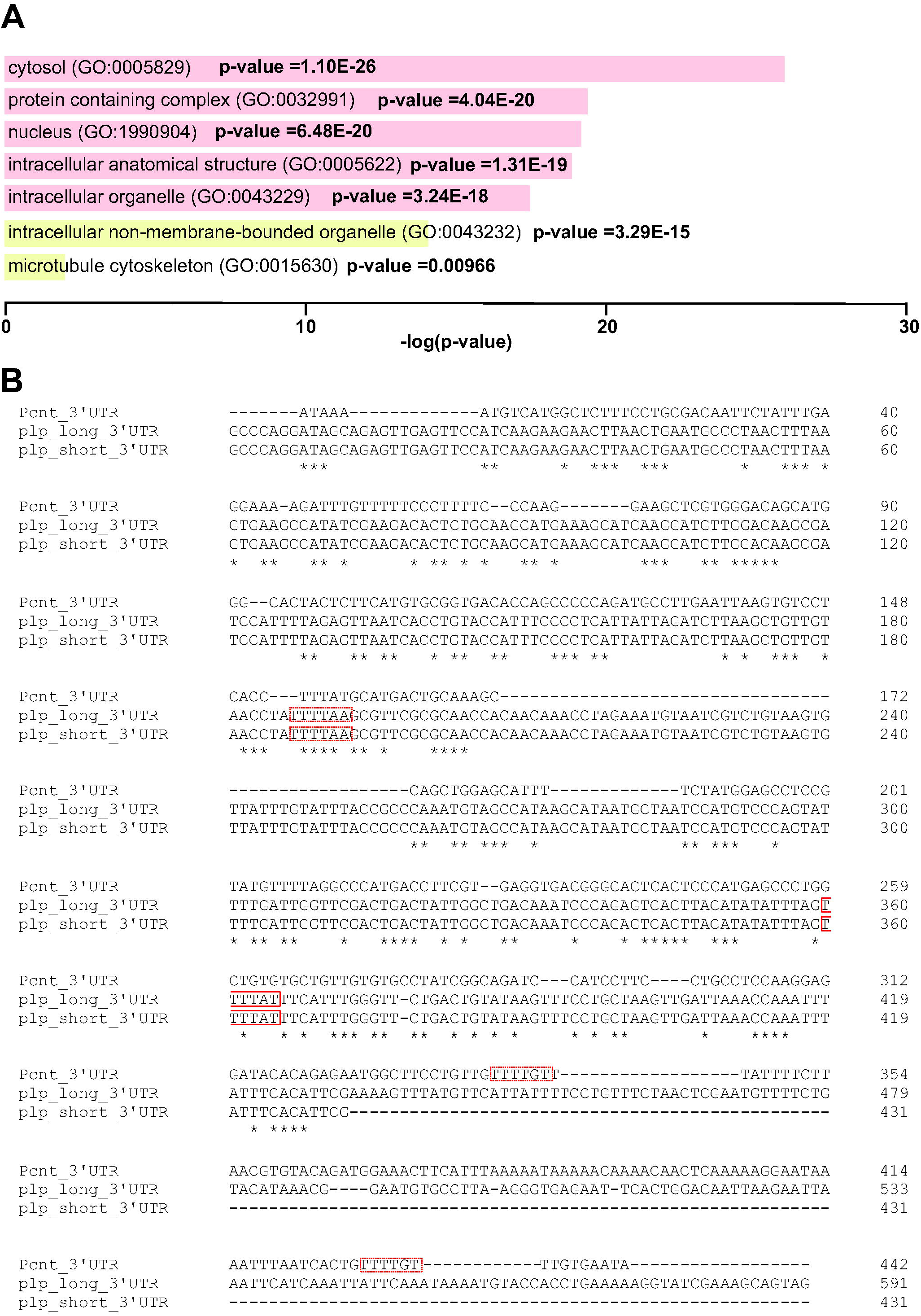
Common mRNA targets of CPEB1 and Orb. (A) Bar graphs show the top 5 most significant cellular component GO terms (pink) and two centrosome-related components (yellow) identified from common mRNA targets of CPEB1 and Orb. Adjusted p-values are displayed. See Supplemental Table 2 for a list of overlapping mRNA targets from the two centrosome-related terms. (B) Alignment of *Drosophila melanogaster plp* and human *PCNT* 3’UTRs. Identical nucleotides are indicated with asterisks. Consensus (red solid boxes) and non-consensus (red dashed boxes) CPE motifs are indicated.

**Figure S3.**
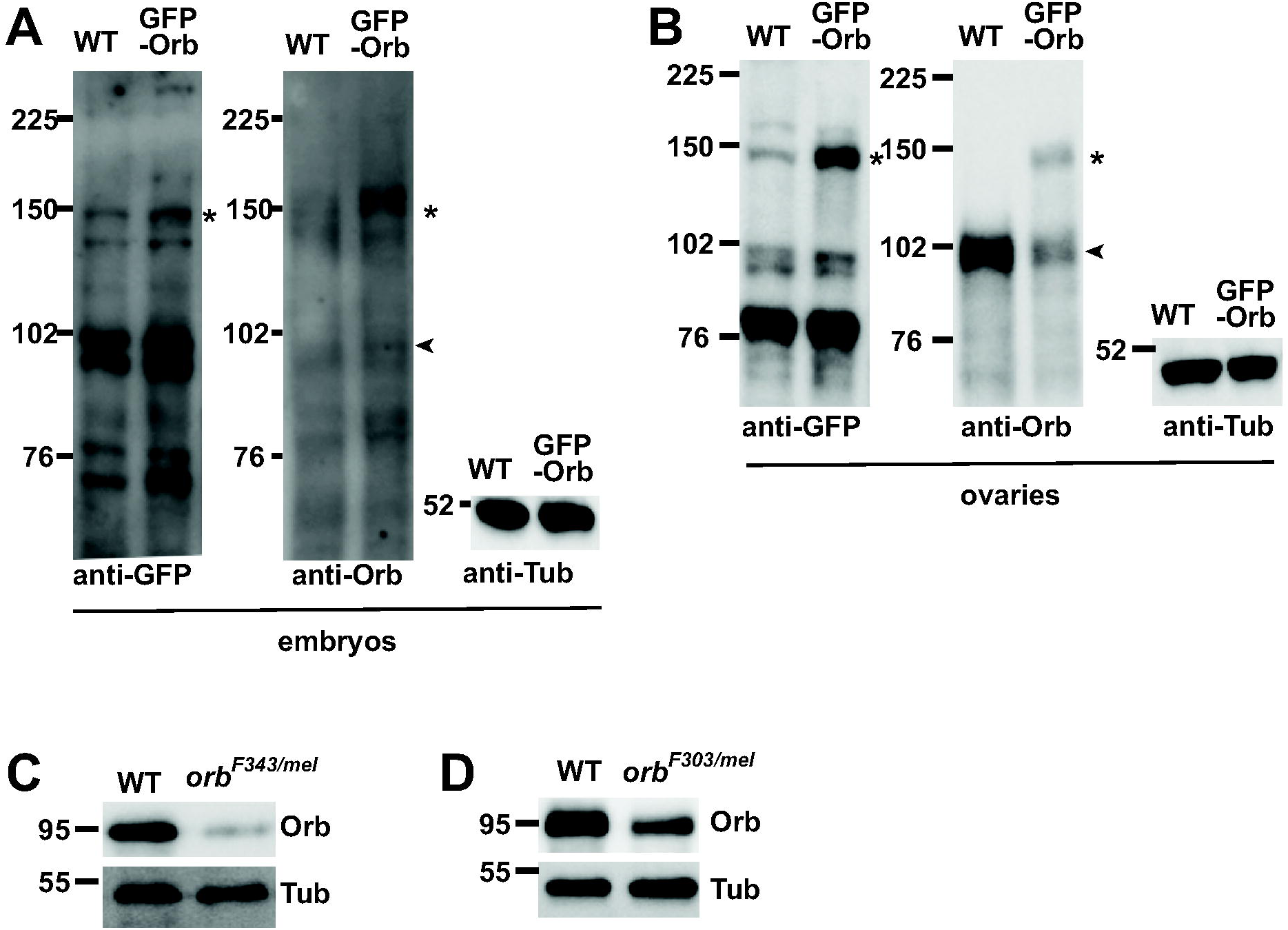
Orb protein expression in ovaries and embryos. (A) Immunoblots of 0-2 hr embryonic extracts from the indicated genotypes probed with anti-GFP, -Orb, and -β-tubulin antibodies. GFP-Orb is expected to migrate near 150 kDa (asterisks), while endogenous Orb protein migrates near 102 kDa (arrowhead). (B) Immunoblots of 2-4 days ovarian extracts from the indicated genotypes stained with anti-GFP, -Orb, and -β-tubulin antibodies. Exogeneous GFP-Orb and endogenous Orb are marked by asterisks and arrowhead, respectively. (C-D) Immunoblots show Orb protein levels from 2-4 day ovarian extracts from WT and *orb* transheterozygous mutants. Uncropped blots are available to view on Figshare: https://doi.org/10.6084/m9.figshare.16900429

**Figure S4.**
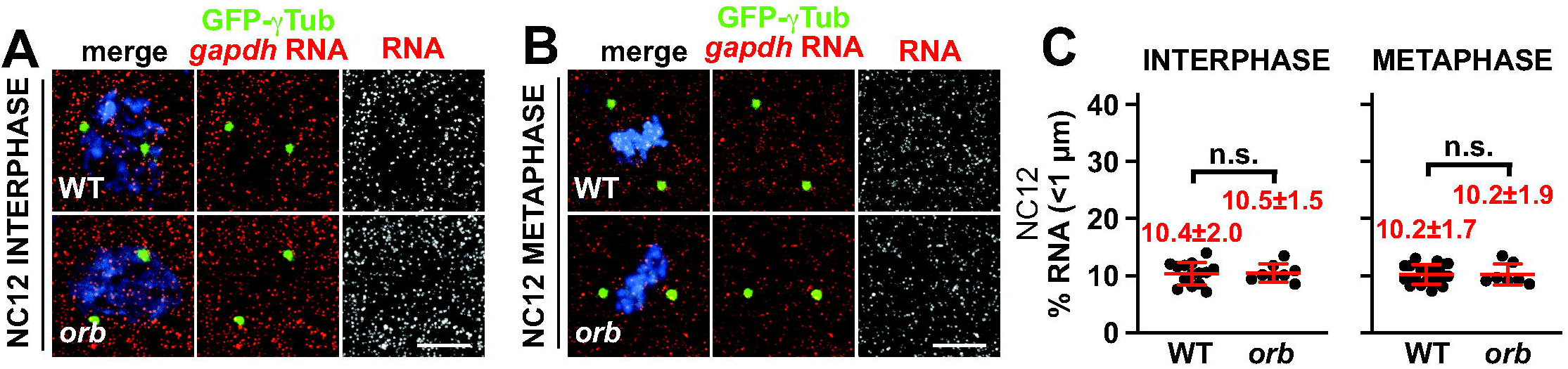
*gapdh* mRNA localization is not affected by *orb* depletion. Images show maximum intensity projections of control and *orb^F343^*/*orb^mel^* mutant NC12 (A) interphase or (B) metaphase embryos expressing *GFP-γTub* (green) and labeled with *gapdh* smFISH probes (red). (C) Quantification of *gapdh* mRNA localization within 1 μm from the centrosome surface in the indicated genotypes. Supplemental Table 1 lists the number of embryos, centrosomes, and RNA objects quantified per condition. n.s, not significant by unpaired t-test. Bars: 5 μm.

**Figure S5.**
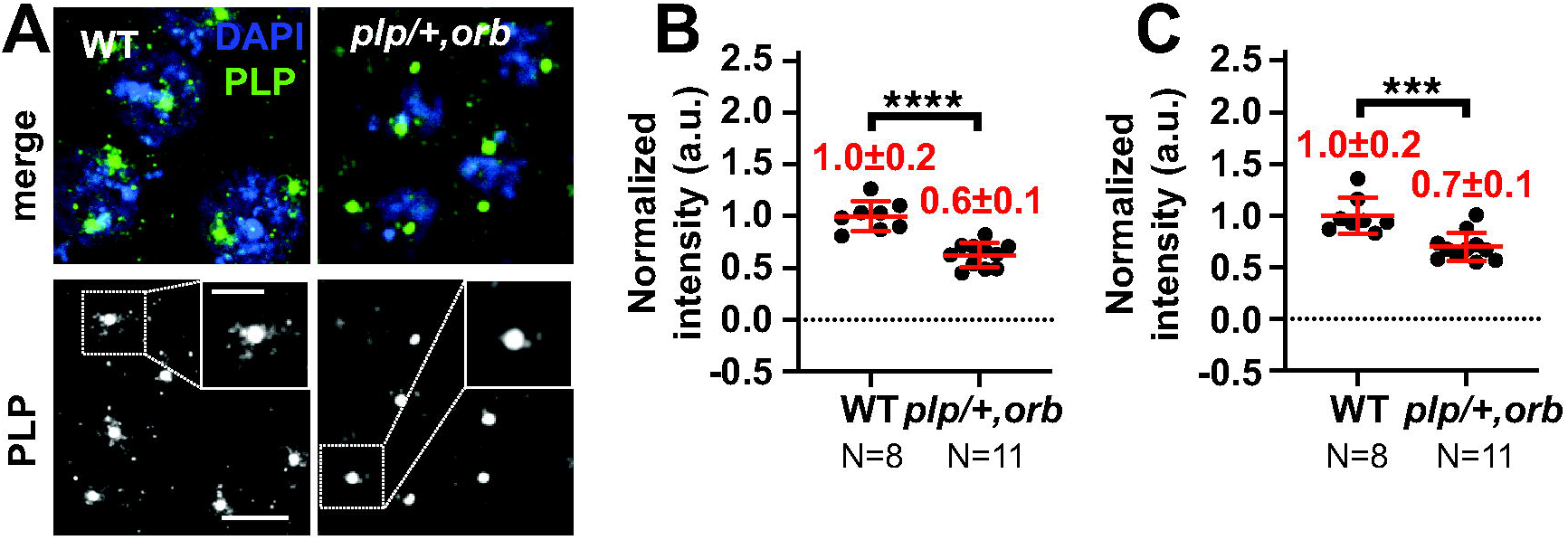
Validation of PLP intensity measurements. (A) Maximum-intensity projections of NC 13 embryos of the indicated genotypes stained with anti-PLP antibodies (green). Quantifications show results from (B) manual quantification of PLP intensity within 2 μm of the centrosome using Fiji vs. (C) a Python-based pipeline, as described in *Materials and Methods*. Each dot represents the average PLP intensity from 10 centrosomes randomly selected from a single embryo. Mean ± S.D. are displayed. \*\*\**p*<0.001 or \*\*\*\**p*<0.0001 by unpaired t-test. *plp*/+,*orb* mutant: *plp^2172^*/+, *orb^F343^*/*orb^mel^*. Bars: 5 μm; 2 μm (insets).

**Figure S6.**
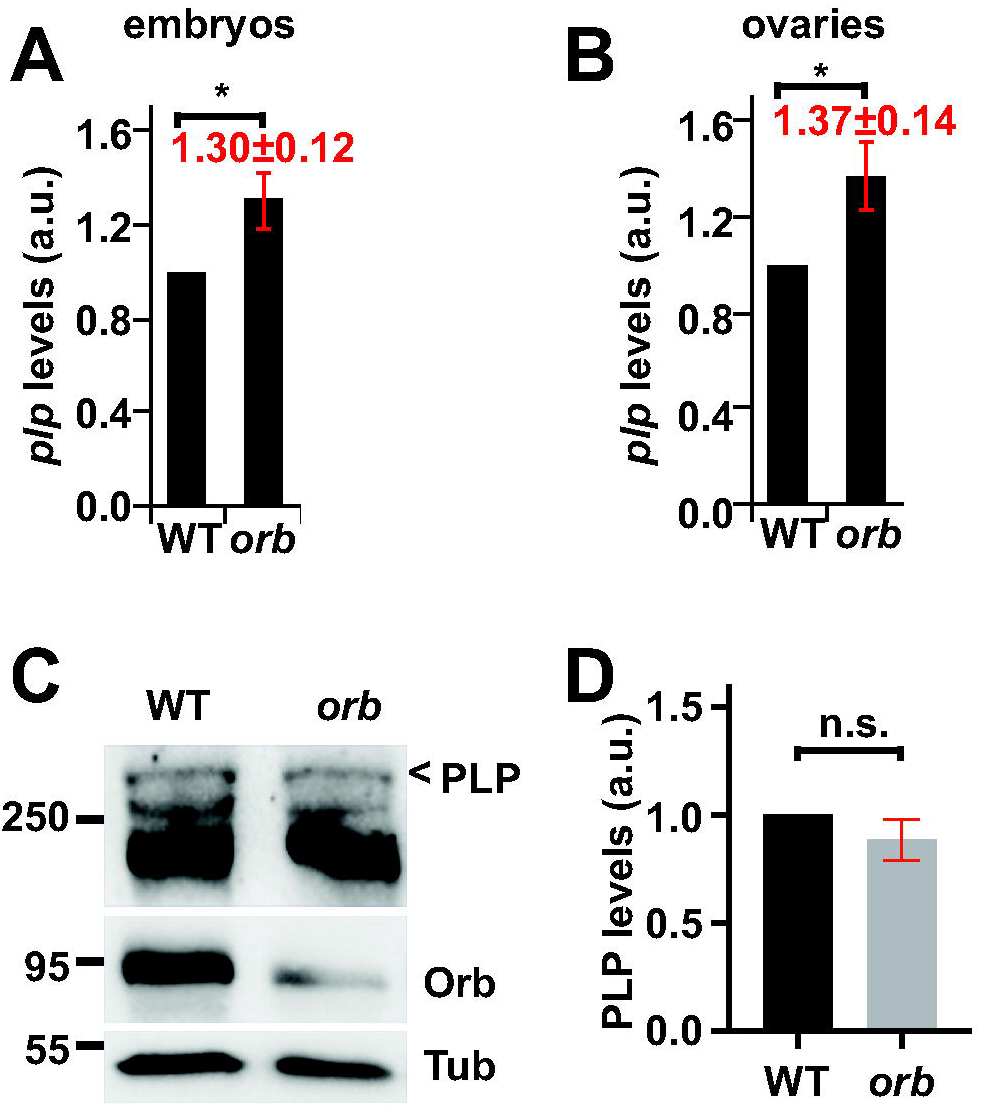
Orb contributes to *plp* mRNA stability. Relative *plp* mRNA levels were examined using qRT-PCR from 0-2 hr embryos (A) or 2–4 day ovarian extracts (B) from the indicated genotypes and normalized to WT. *orb* mutant: *orb^F343^*/*orb^mel^*. (C) Immunoblots show relative PLP protein levels from 2–4 day ovarian extracts. Top PLP bands (carrot) were quantified, as in (D). (D) Quantification of normalized relative PLP levels from three biological replicates. n.s, not significant; Mean ± S.D. are displayed. *orb* mutant: *orb^F343^*/*orb^mel^*. \**p*<0.05 by unpaired t-test. Uncropped blots are available to view on Figshare: https://doi.org/10.6084/m9.figshare.16900423.v1

## Supplemental Materials

**Table S1. Summary of the quantification of RNA localization to centrosomes.**

For every image quantified, the following parameters are listed: A) Figure, B) Genotype, C) mRNA analyzed, D) NC stage of embryos, E) cell cycle phase of embryos, F) (n) embryos quantified, G) (n) centrosomes segmented, H) total number (n) of mRNA objects detected, I) all mRNA objects (n) within 1 μm from the centrosome surface, J) total number (n) of single molecules of mRNA after single molecule normalization, and K) number of single molecules of mRNA within 1 μm from the centrosome surface. The values from Column K are displayed in the figures.

**Table S2. Common mRNA targets of CPEB1 and Orb.**

Three lists of overlapping mRNA targets shared between Orb and CPEB1 are provided: 1) all common genes, 2) common genes within GO-term ‘microtubule cytoskeleton,’ and 3) common genes within GO-term ‘non-membrane organelle.’ *PCNT*/*plp* are highlighted.

**Supplemental Video 1. GFP-γTub and H2A-RFP from NC 12 interphase to NC 14 interphase in a control embryo.** Live control *Drosophila* embryo expressing *GFP-γTub* (green) and *H2A-RFP* (red). Frames were captured at 20 s intervals over 31 min 40 s using a 1 μm z-step size over 14 μm total depth. Video displayed using a playback speed of 6 FPS (frame per second). Bar: 10 μm.

**Supplemental Video 2. GFP-γTub and H2A-RFP from NC 12 interphase to NC 14 interphase in an *orb* mutant embryo.** *Drosophila* embryo expressing *GFP-γTub* (green) and *H2A-RFP* (red). Frames were captured every 20 s for 29 min 40 s using a 1 μm z-step size over 14 μm total depth. Images were captured at 1 μm z-intervals over a 10–15 μm volume at 20 s intervals. Video displayed with a playback speed of 6 FPS. *orb* mutant: *orb^F303^*/*orb^mel^*. Bar: 10 μm.

**Supplemental Video 3. GFP-γTub and H2A-RFP from NC 12 interphase to NC 14 interphase in a *PLP-GFP*/+; *orb* mutant embryo.** *Drosophila* embryo expressing *GFP-γTub* (green) and *H2A-RFP* (red). Frames were captured every 22 s for 30 min 04 s using a 1 μm z-step size over 15 μm total depth. Video is displayed using a playback speed of 6 FPS. *PLP- GFP/+*; *orb* mutant: *PLP-GFP/+*; *orb^F303^*/*orb^mel^*. Bar: 10 μm.

